# Transcriptome profiling of mouse samples using nanopore sequencing of cDNA and RNA molecules

**DOI:** 10.1101/575142

**Authors:** Camille Sessegolo, Corinne Cruaud, Corinne Da Silva, Audric Cologne, Marion Dubarry, Thomas Derrien, Vincent Lacroix, Jean-Marc Aury

## Abstract

Our vision of DNA transcription and splicing has changed dramatically with the introduction of short-read sequencing. These high-throughput sequencing technologies promised to unravel the complexity of any transcriptome. Generally gene expression levels are well-captured using these technologies, but there are still remaining caveats due to the limited read length and the fact that RNA molecules had to be reverse transcribed before sequencing. Oxford Nanopore Technologies has recently launched a portable sequencer which offers the possibility of sequencing long reads and most importantly RNA molecules. Here we generated a full mouse transcriptome from brain and liver using the Oxford Nanopore device. As a comparison, we sequenced RNA (RNA-Seq) and cDNA (cDNA-Seq) molecules using both long and short reads technologies and tested the TeloPrime preparation kit, dedicated to the enrichment of full-length transcripts. Using spike-in data, we confirmed that expression levels are efficiently captured by cDNA-Seq using short reads. More importantly, Oxford Nanopore RNA-Seq tends to be more efficient, while cDNA-Seq appears to be more biased. We further show that the cDNA library preparation of the Nanopore protocol induces read truncation for transcripts containing internal runs of T’s. This bias is marked for runs of at least 15 T’s, but is already detectable for runs of at least 9 T’s and therefore concerns more than 20% of expressed transcripts in mouse brain and liver. Finally, we outline that bioinformatics challenges remain ahead for quantifying at the transcript level, especially when reads are not full-length. Accurate quantification of repeat-associated genes such as processed pseudogenes also remains difficult, and we show that current mapping protocols which map reads to the genome largely over-estimate their expression, at the expense of their parent gene. The entire dataset is available from http://www.genoscope.cns.fr/externe/ONT_mouse_RNA.

## Introduction

To date our knowledge of DNA transcription is brought by the sequencing of RNA molecules which have been first reverse transcribed (RT). This RT step is prone to skew the transcriptional landscape of a given cell and erase base modifications. The sequencing of these RT-libraries, that we suggest to call cDNA-Seq, has became popular with the introduction of the short-read sequencing technologies [28], [19]. Recently, the Oxford Nanopore Technologies (ONT) company commercially released a portable sequencer which is able to sequence very long DNA fragments [7] and enable now the sequencing of complex genomes ([9], [3] and [23]). Moreover this device (namely MinION) is also able to sequence native RNA molecules [8] representing the first opportunity to generate genuine RNA-Seq data.

Furthermore, even if short-read technologies offer a deep sequencing and were helpful to understand the transcriptome complexity and to improve the detection of rare transcripts, they still present some limitations. Indeed, read length is a key point to address complex regions of a studied transcriptome. Depending on the evolutionary history of a given genome, recent paralogous genes can lead to ambiguous alignment when using short reads. In a same way, processed pseudogenes generated by the retrotranscription of RNAs back into the genomic DNA are challenging to quantify using short reads. In addition to sequencing technologies and bioinformatics methods, preparation protocols have a significant impact on the final result as they can incorporate specific biases [1],[27]. The generation of data rely on a high number of molecular and computational steps which evolve at a fast pace. These changes in the protocol generally modified the appearance of the data. As an example, data produced with protocols based on oligo-dT or random primers in the RT step show differences in how they cover transcripts[1].

## Results

### Experimental design

Here we produce a complete transcriptome dataset, containing both cDNA-Seq and RNA-Seq, using the Illumina and Nanopore technologies. RNAs were sampled from brain and liver tissues of mice and were mixed with Lexogen’s Spike-In RNA Variants (SIRVs) as a control for quantification of RNAs. We follow the protocols recommended by the manufacturers to generate the three following datasets on each tissue: Illumina cDNA-Seq, Nanopore cDNA-Seq and Nanopore RNA-Seq. The first was sequenced using the Illumina platform (TruSeq_SR) and the last two using the MinION device (PCS108_LR and RNA001_LR). From the brain tissue, we generated biological (two brain RNA samples, C1 and C2) and technical replicates (R1 and R2) for the three datasets (Figure 1). Additionally, the second was also sequenced using the Illumina platform (PCS108_SR). This enables us to clarify which differences are due to the preparation protocol and which are due to the sequencing platform in itself. Moreover, we generated a Lexogen’s TeloPrime library on both tissues (TELO_LR), this preparation kit is an all-in-one protocol for generating full-length cDNA from total RNA (Figure 1 and Tables 1 and 2).

**Figure 1:**
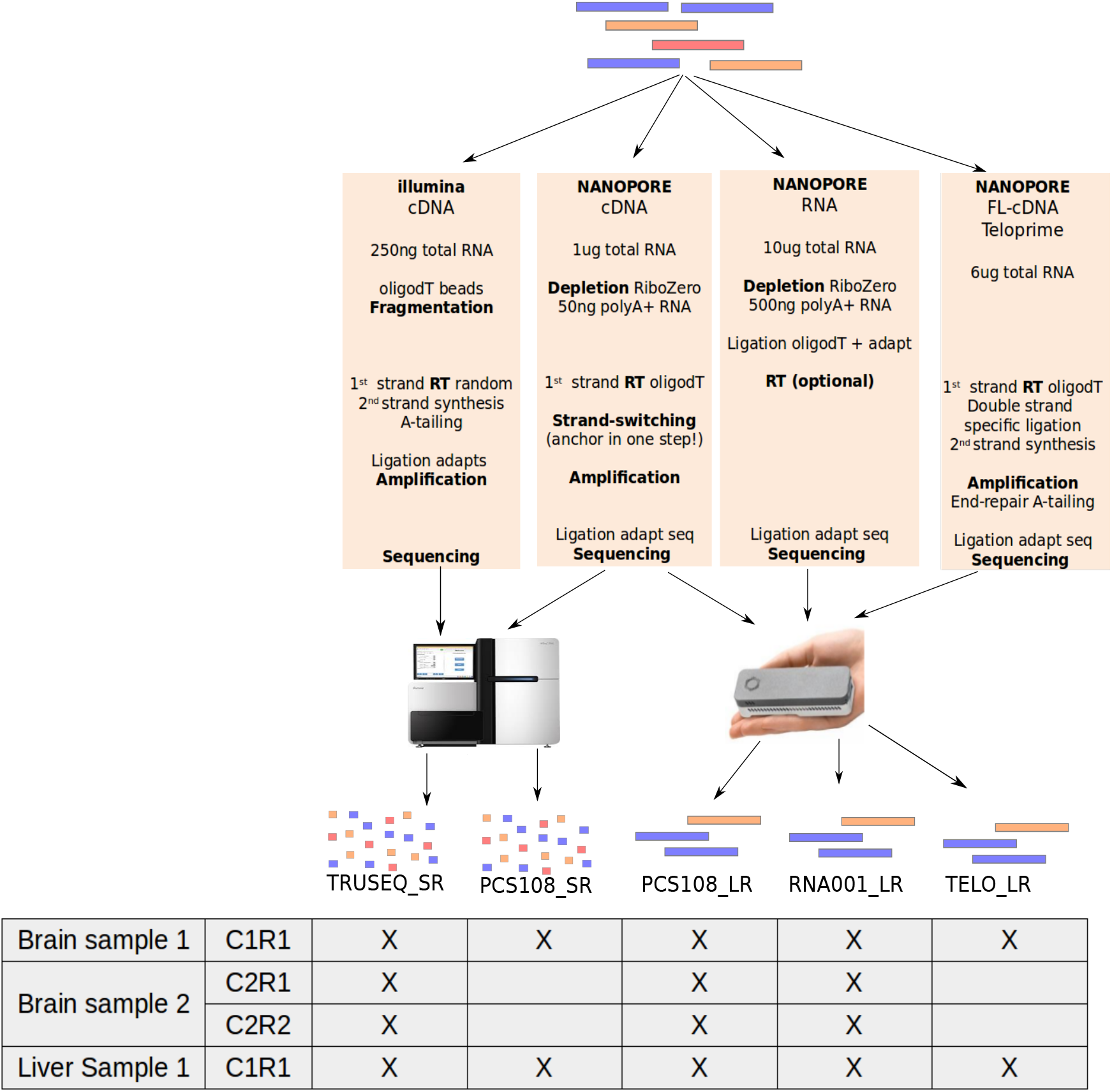
Experimental design. Five protocols have been used on each tissue. Two were based on short-reads with the TruSeq protocol (TRUSEQ_SR) and the ONT library preparation (PCS108_SR) and the three others were based on long-reads with the ONT cDNA-Seq protocol (PCS108_LR), the ONT RNA-Seq protocol (RNA001_LR) and the Teloprime protocol (TELO_LR). (RT : Reverse Transcription). For the brain, two biological replicates, C1 and C2, have been generated and two technical replicates, R1 and R2, have been generated for the second biological replicate. For the first biological replicate all the five protocols were used whereas the TRUSEQ_SR, the PCS108_LR and the RNA001_LR were used for the second biological replicates.

**Table 1:** Standard metrics of the generated ONT datasets for both tissues : brain and liver.

|  | PCS108_LR |  |  | RNA001_LR |  |  | TELO_LR |
| --- | --- | --- | --- | --- | --- | --- | --- |
| RNA sample | C1 | C2 |  | C1 | C2 |  | C1 |
| Replicate | R1 | R1 | R2 | R1 | R1 | R2 | R1 |
|  | Brain |  |  |  |  |  |  |
| Number of reads | 1,267,830 | 5,834,882 | 3,003,844 | 571,098 | 364,041 | 210,654 | 1,691,454 |
| Cumulative size (Gb) | 1.30 | 7.03 | 3.28 | 0.43 | 0.38 | 0.20 | 1.31 |
| Average Size (bp) | 1,028.94 | 1,204.54 | 1,091.85 | 758.05 | 1,032.10 | 957.17 | 775.77 |
| N50 (bp) | 1,283 | 1,749 | 1,591 | 1,357 | 1,492 | 1,417 | 896 |
| Number reads >1Kb | 522,422 | 2,869,633 | 1,339,489 | 154,735 | 141,970 | 73,232 | 389,468 |
| Accession number | ERX2695238 | ERX3387950 | ERX3387952 | ERX2695236 | ERX3387949 | ERX3387951 | ERX2850744 |
|  | Liver |  |  |  |  |  |  |
| Number of reads | 3,043,572 | - | - | 418,102 | - | - | 2,668,975 |
| Cumulative size (Gb) | 3.30 | - | - | 0.34 | - | - | 2.64 |
| Average Size (bp) | 1,083.85 | - | - | 823.30 | - | - | 989.85 |
| N50 (bp) | 1,264 | - | - | 1,153 | - | - | 1,116 |
| Number of reads >1Kb | 1,218,569 | - | - | 117,047 | - | - | 884,008 |
| Accession number | ERX2695243 | - | - | ERX2695240 | - | - | ERX2850745 |

**Table 2:** Standard metrics of the Illumina datasets for both tissues : brain and liver.

|  | TruSeq_SR |  |  | PCS108_SR |
| --- | --- | --- | --- | --- |
| RNA sample | C1 | C2 |  | C1 |
| Replicate | R1 | R1 | R2 | R1 |
|  | Brain |  |  |  |
| Number of reads | 53,128,934 | 41,562,993 | 45,719,216 | 153,610,181 |
| Cumulative size (Gb) | 15.42 | 12.36 | 13.60 | 45.88 |
| Read Size (bp) | 151 | 151 | 151 | 151 |
| Accession number | ERX2695239 | ERX3387947 | ERX3387948 | ERX2695237 |
|  | Liver |  |  |  |
| Number of reads | 49,270,153 | - | - | 178,019,939 |
| Cumulative size (Gb) | 14.16 | - | - | 53.43 |
| Read Size (bp) | 151 | - | - | 151 |
| Accession number | ERX2695241 | - | - | ERX2695242 |

### Spliced alignment and error rate

The error rate of ONT reads is still around 10% and complicates the precise detection of splice sites. Mouse splice sites are often canonical, as observed when aligning reference annotation (coding genes from Ensembl 94) using BLAT, 98.5% of introns were GT-AG. Here minimap2 was able to detect only 80.7% of GT-AG introns when ONT RNA-Seq reads were used as input. Interestingly, the proportion of cannonical splice sites is lower when using ONT cDNA-Seq (67.7%). In fact ONT RNA-Seq reads are strand-specific which is of high value for the alignment and splice site detection. When using high quality sequences (coding genes from Ensembl 94) instead of reads, minimap2 retrieved 96.4% of GT-AG introns. These results show that the detection of splice sites using long but noisy reads is challenging and that dedicated aligners still need some improvements.

### General comparison of sequencing technologies

RNA-Seq is a powerful method that provides a quantitative view of the transcriptome with the number of sequenced fragments being a key point to thoroughly capture the expression of genes. The Illumina technology is able to generate billions of short tags, and unsurprisingly allows to access a largest number of genes/transcripts. However with the same number of reads, both Illumina and Nanopore technology are able to uncover the same number of transcripts (Figure 2).

**Figure 2:**
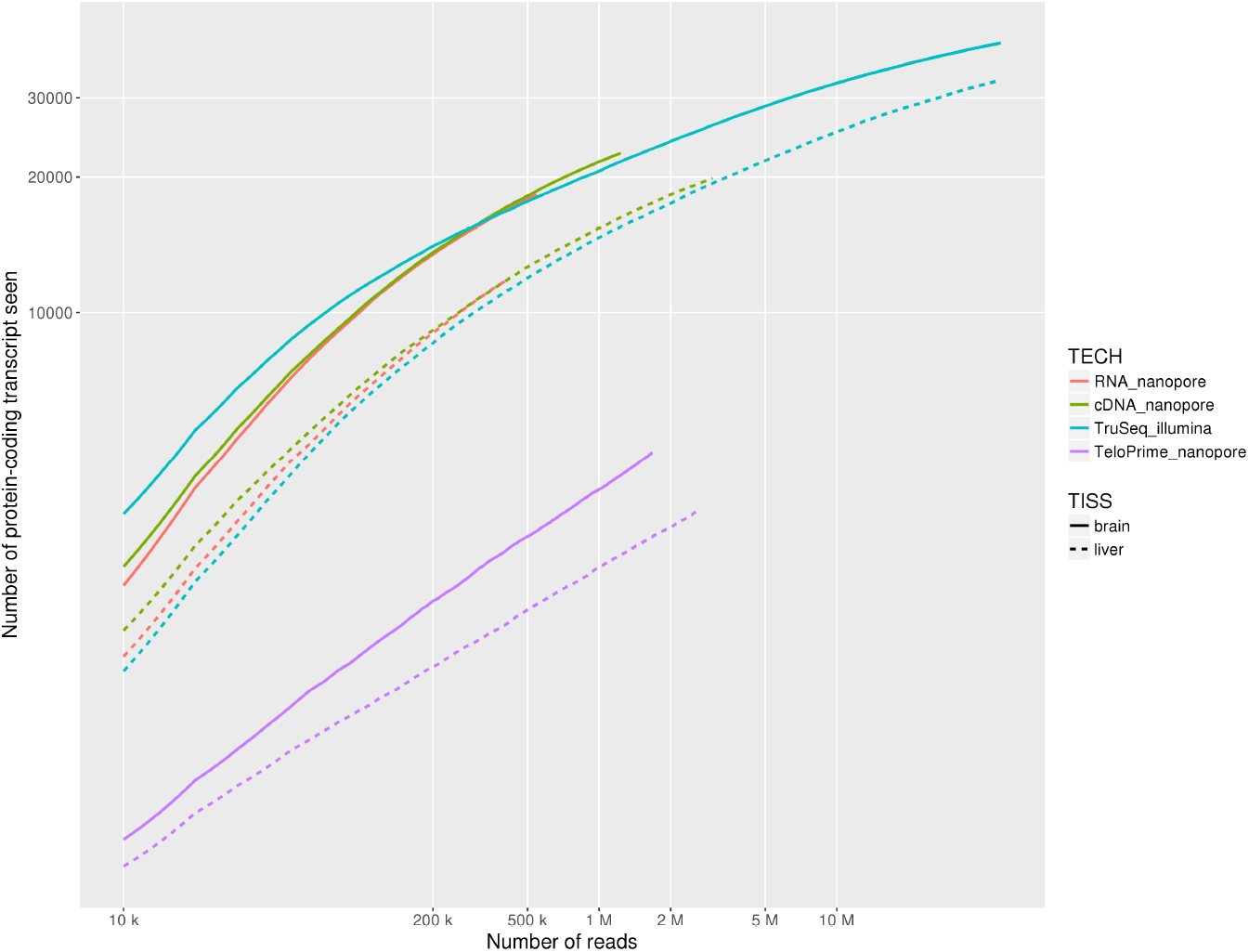
Saturation curve. Number of protein-coding transcripts seen by each technology at various sequencing depth. Solid and dashed lines correspond respectively to brain and liver samples.

Long reads sequencing offers the possibility to capture full-length RNAs. When looking at single isoform genes, we found that in average reads cover between 61 and 74% of the messenger RNAs (Table 3). But even though this horizontal coverage is quite high, the proportion of reads that covered more than 80% of the transcript remains low (near 55% except for cDNA-Seq of the brain sample). RNA degradation can obviously explain a proportion of these fragmented reads, and it has been shown more recently that a software artifact may truncate reads (around 20%) during the base-calling process [4].

**Table 3:** Long reads coverage of single-isoform genes.

|  | PCS108_LR |  |  | RNA001_LR |  |  | TELO_LR |
| --- | --- | --- | --- | --- | --- | --- | --- |
| RNA sample | C1 | C2 |  | C1 | C2 |  | C1 |
| Replicate | R1 | R1 | R2 | R1 | R1 | R2 | R1 |
|  | Brain |  |  |  |  |  |  |
| # of mapped reads | 99,192 | 389,007 | 203,622 | 45,485 | 48,128 | 29,719 | 200,884 |
| Avg coverage | 61.18% | 62.18% | 62.10% | 70.97% | 74.25% | 76.52% | 76.37% |
| Median coverage | 64.62% | 62.11% | 61.59% | 83.84% | 89.34% | 92.36% | 84.65% |
| Full-length reads (>80%) | 40.25% | 41.08% | 40.73% | 53.31% | 57.19% | 60.61% | 60.18% |
|  | Liver |  |  |  |  |  |  |
| # of mapped reads | 344,362 | - | - | 48,032 | - | - | 381,005 |
| Avg coverage | 73.86% | - | - | 73.48% | - | - | 79.09% |
| Median coverage | 83.24% | - | - | 84.57% | - | - | 88.23% |
| Full-length reads (>80%) | 55.63% | - | - | 54.77% | - | - | 60.02% |

### Improving the proportion of full-length reads

The proportion of full-length transcripts can be improved by using a dedicated library preparation protocol. Here we tested the TeloPrime amplification kit, commercialized by the Lexogen company. This protocol is selective for full-length RNA molecules that are both capped and polyadenylated. Using this protocol we were able to slightly improve the proportion of full-length reads (which covered at least 80% of a given transcript, Table 3). However, in return, we captured a lower number of genes, lowering the interest of such a protocol in most applications (Figure 2).

### Sequencing biases of transcripts containing internal runs of poly(T)

Since cDNA synthesis is initiated with an anchored poly-dT primer (poly-TVN), a relevant question is whether transcripts containing internal runs of poly(A) or poly(T) are correctly sequenced. We computed the relative coverage for each transcript upstream and downstream internal runs of poly(A) or poly(T) (see Methods), and we find that using cDNA-Seq, cDNA molecules stemming from such transcripts are often truncated. This bias is detectable for runs of poly(A) (Figure 3b) but is much stronger for runs of poly(T) (Figure 3a). While the first situation could be caused by internal poly(T) priming during first strand cDNA synthesis and therefore result in 3’-truncation of the cDNA, the second situation could occur during 2nd strand cDNA synthesis and result in 5’-truncation of the cDNA (as sketched in Figure 3c). As an example, the *Set* gene contains an internal run of 20 T’s and ONT cDNA-Seq reads are systematically interrupted at this location (Figure 3c, tracks 2 and 3), while this is not the case for Illumina Truseq (Figure 3d, track 4) and ONT RNA-Seq (Figure 3d, track 1). We find that the magnitude of the bias is associated to the length of the internal run of poly(T) (Supplementary Figure 1). The bias is very pronounced for transcripts containing at least 15 T’s, but it is already detectable for transcripts containing at least 9 T’s. This bias has remained unreported so far, but it is also present in other published Nanopore dataset [30] (Supplementary figure 2). It however concerns a large fraction of expressed transcripts. Indeed, transcripts containing at least 9 T’s correspond to 27% of transcripts expressed with at least one read in mouse brain (resp. 20% in mouse liver). In human GM12878 cell line, this proportion is 16%. Importantly, the bias not only affects read length, but also transcript quantification. Indeed, cDNA-Seq reads from these transcripts are not only shorter, they are also more numerous. As an example, the *set* gene is covered by 497 truncated cDNA reads and 22 full-length RNAseq reads (Figure 3d). More generally, in mouse brain, 35% of cDNA-Seq reads map to transcripts with at least 9 T’s, compared to 14% of RNA-Seq reads. This suggests that the abundance of these transcripts is over-estimated when using cDNA-Seq, at the expense of the other transcripts.

**Figure 3:**
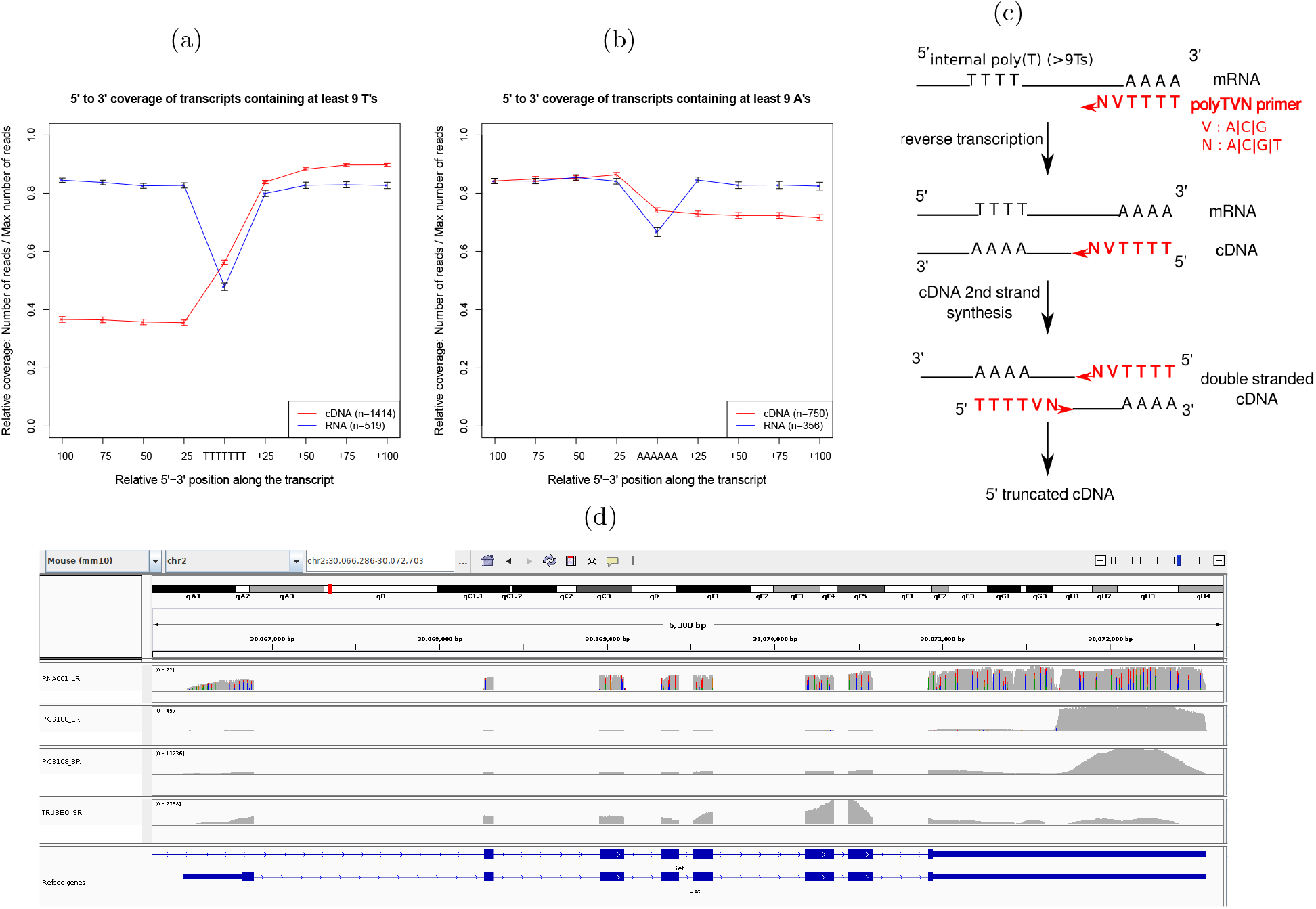
Truncated reads. (a) Relative coverage of transcripts for the ONT cDNA-Seq dataset and the ONT RNA-Seq dataset for transcripts covered by at least 10 reads around a poly(T). With the ONT cDNA-Seq dataset, transcripts containing internal runs of at least 9 T’s are less covered in 5’. The coverage deficit observed in the ONT RNA-seq dataset is due to indel sequencing errors associated to homopolymers. (b) Relative coverage of transcripts for the ONT cDNA-Seq dataset and the ONT RNA-Seq dataset for transcripts covered by at least 10 reads around a poly(A). Using the ONT cDNA-Seq dataset, transcripts containing stretches of at least 9 A’s are less covered in 3’. Again, the coverage deficit observed in the ONT RNA-seq dataset is due to indel sequencing errors associated to homopolymers. (c) Mechanism explaining why internal runs of T’s are causing 5’ truncated reads. The PolyTVN primer binds to the internal run of poly(A) of the cDNA so that the second cDNA strand is 5’ truncated. (d) Example of a gene named *Set* visualized with IGV. Truncated reads are in tracks 2 (ONT cDNA-Seq) and 3 (Illumina, Nanopore protocol). Non-truncated reads are in tracks 1 (ONT RNA-Seq) and 4 (Illumina Truseq). The region where the truncation occurs is a poly(T).

### Evaluation of the accuracy of the gene expression quantification using spike-in data

In order to assess which protocol was best to quantify gene expression, we analyzed the 67 spike-ins contained in the brain datasets. Since we exactly know which transcripts are present in the sample, the quantification is rather straightforward. We aligned reads to the reference transcriptome, used RSEM for short reads, and counted the number of primary alignments for long reads (see Methods). The best quantifications were obtained for the ONT RNA-Seq (Spearman *ρ* =0.86, Pearson *r*=0.85) and Illumina TruSeq (*ρ* =0.81, *r*=0.82) protocols (Figure 4). In contrast, cDNA-Seq (sequenced using Illumina or ONT) produced more imprecise quantifications (*ρ* =0.54, *r* = 0.57 and *ρ* =0.6, *r* = 0.50). Importantly, we obtained very similar results on all our three replicates (Supplementary Figure 3, 4), with ONT RNA-Seq consistently exhibiting the higher correlation with true quantifications. The use of salmon either for short or long reads, as was done in [25] did not change our results. We then wanted to test if the number of ONT RNA-Seq reads was indeed a better predictor of the true transcript quantification, than the number of cDNA-Seq reads or the TPM measure derived from Illumina. Using 30 fold crossvalidations, we found that the mean square error was 1.56 ± 0.01 for ONT cDNA-Seq, 1.38 ± 0.03 for Illumina and 0.77 ± 0.01 for ONT RNA-Seq. When inspecting the errors made for each SIRV, we noticed that SIRV311 was particularly poorly predicted by all methods (possibly because it is only 191nt long which makes it the shortest SIRV), and in particular by Illumina TruSeq. When removing it from the dataset, we obtained *ms_illu_* ∈ [0.623; 0.634], *ms_rna_* ∈ [0.483; 0.498], *ms_cDNA_* ∈ [1.44; 1.47] which highlights that ONT RNA-Seq yields significantly better quantifications than Illumina TruSeq and ONT cDNA-Seq. Although the magnitude of the difference with Illumina TruSeq is small, we found it to be reproducible. We could further show that, for each technology, the errors made for each SIRV were reproducible across replicates (Supplementary Figure 5) meaning that a transcript whose expression is over-estimated with one technology is consistently over-estimated with the same technology.

**Figure 4:**
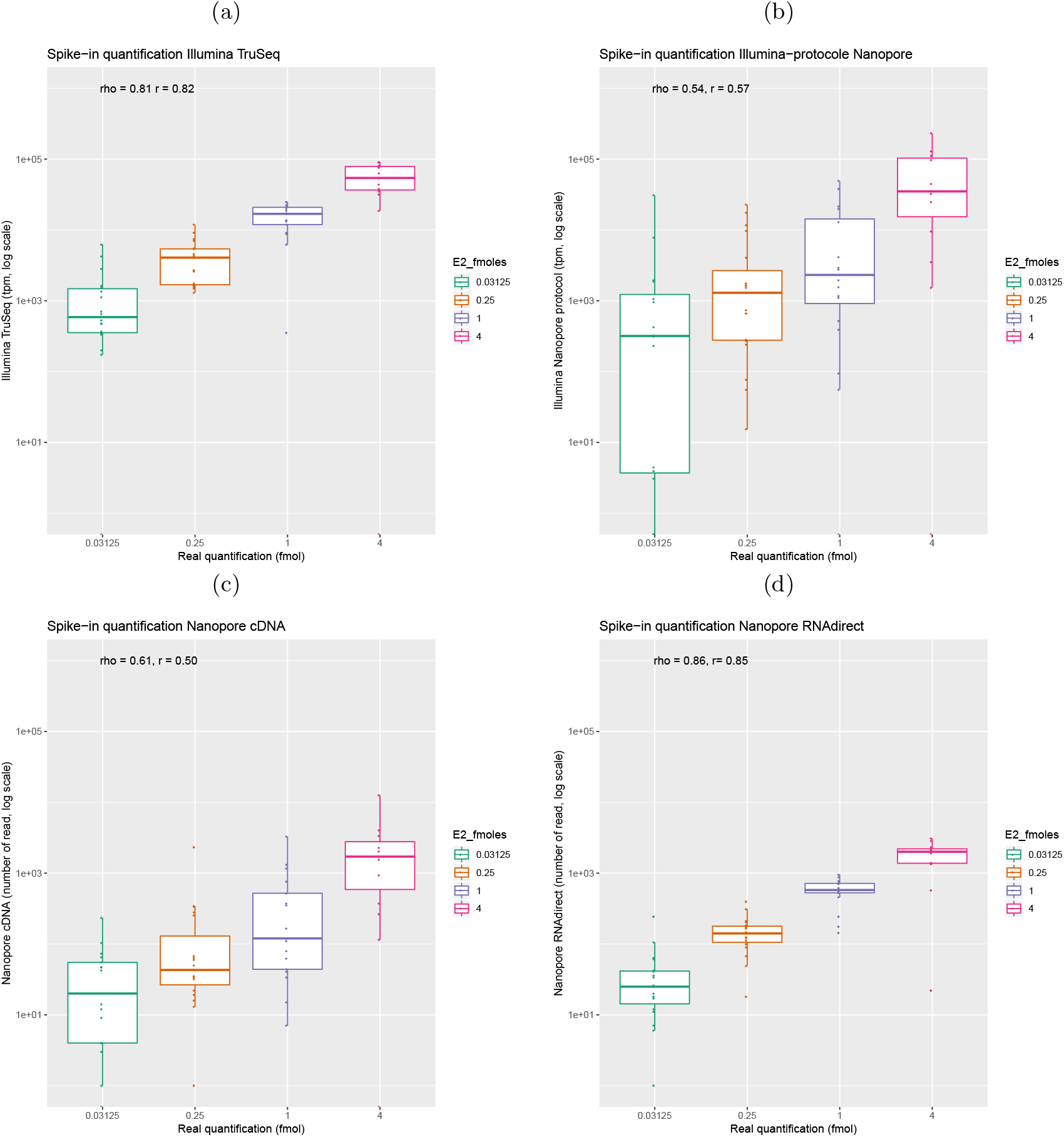
Evaluation of quantification using the SIRV E2 spike-in mix. Reads were mapped against the SIRV transcriptome and quantifications were computed at the transcript level. The observed quantifications are correlated with the known theoretical quantifications of the spike in. (a) Correlation obtained for Illumina cDNA-Seq (Spearman’s *ρ* = 0.80, n=327,386). (b) Correlation obtained for Illumina with the ONT cDNA-Seq protocol (Spearman’s *ρ* =0.53, n=7,981,494 reads). (c) Correlation obtained for ONT cDNA-Seq (Spearman’s *ρ* = 0.65, n=42,559 reads). (d) Correlation obtained for ONT RNA-Seq (Spearman’s *ρ* = 0.86, n=35,513 reads).

In order to assess the quality of the quantification in a more realistic context where we do not know which transcripts are present in the sample, we also mapped the reads to a modified set of transcripts corresponding either to an over-annotation or an under-annotation (as provided by Lexogen). In both cases, the correlations were overall poorer than before, but the order was maintained, with ONT RNA-Seq and then Illumina cDNA-Seq being the more reliable protocols (Supplementary Figures 6 and 7).

### Quantification of the expression level of mouse transcripts

Given that with the spike in, the best quantification were obtained with ONT RNA-Seq, we compared the quantifications obtained with this protocol with the ones obtained with the ONT and Illumina cDNA-Seq protocols. Figure 5 summarizes the correlations in terms of transcript quantification of our datasets. Comparing the Illumina cDNA-Seq and the ONT RNA-Seq protocols we obtain a spearman coefficient of correlation *ρ* = 0.51. The correlation is lower in the liver sample (Supplementary Figure 8) probably because of a lower number of RNA-seq reads and a shorter read length.

**Figure 5:**
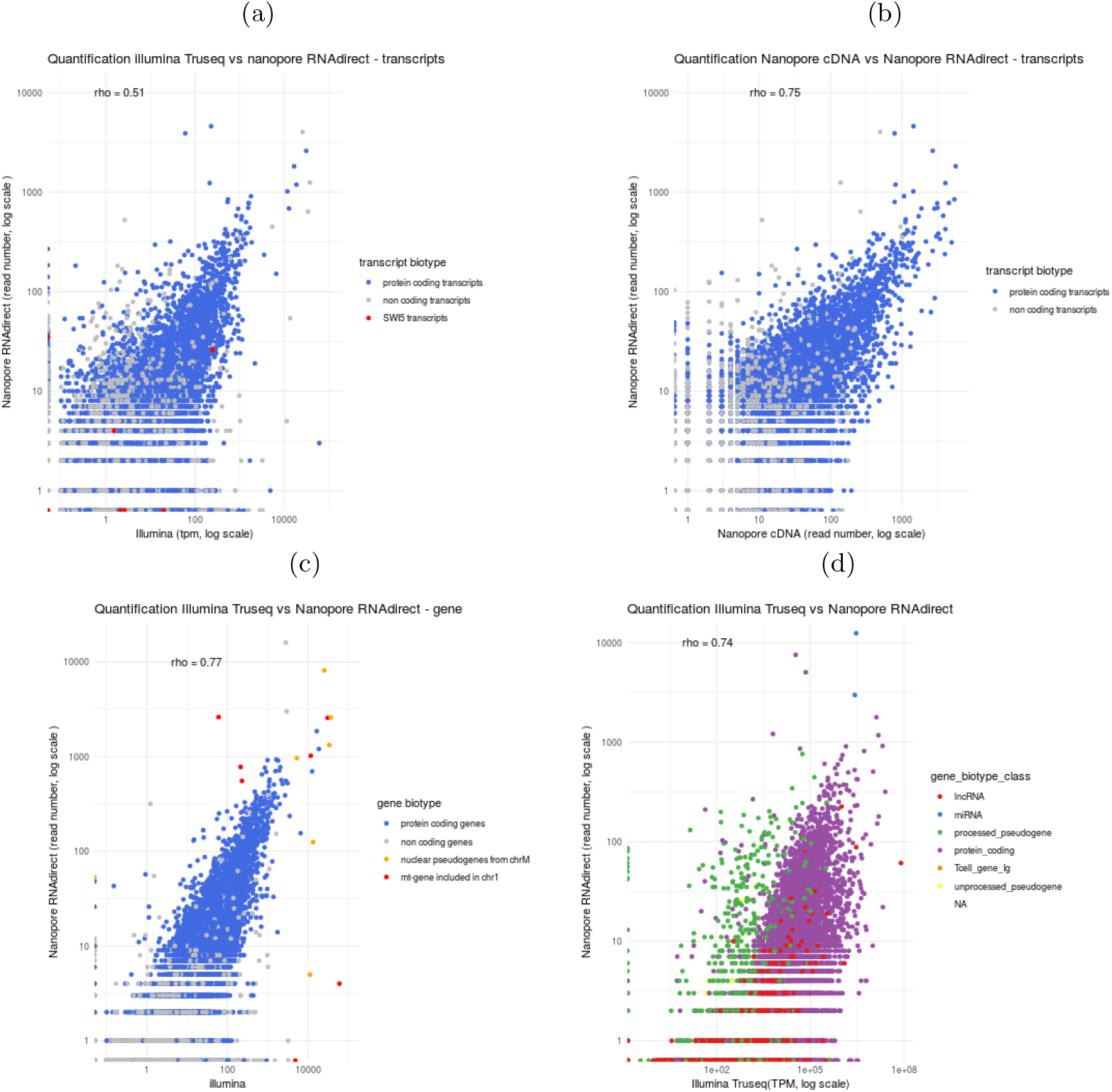
Comparison of quantifications. Transcripts or genes annotated as protein-coding are in blue. Spearman’s *ρ* has been computed for all transcripts or genes. (a) Comparison of ONT RNA-Seq and Illumina cDNA-Seq quantifications at the transcript level (Spearman’s *ρ* = 0.51). Red points correspond to the transcripts of the *Swi5* gene. (b) Comparison of ONT RNA-Seq and ONT cDNA-Seq quantifications at the transcript level (Spearman’s *ρ* = 0.75). (c) Comparison of ONT RNA-Seq and Illumina cDNA-Seq quantifications at the gene level (transcript quantification were summed for each gene, Spearman’s *ρ* = 0.77). Red points correspond to pseudogenes located on chromosome 1 within the NUMT, i.e. segment of the mitochondrial genome which has been copied and integrated in the nuclear genome and orange points correspond to the original mitochondrial genes. (d) Reads were mapped against the mouse reference genome and quantifications computed at gene level. We compared the ONT RNA-Seq and the Illumina cDNA-Seq protocols (Spearman’s *ρ* = 0.74). Green points correspond to processed pseudogenes, red points to long non coding RNAs.

Comparing the ONT RNA-Seq and cDNA-Seq quantification, we obtain a higher correlation (*ρ* = 0.75), suggesting that read length strongly influences transcript quantification. Indeed, in the comparison between Illumina cDNA-Seq and ONT RNA-Seq dataset, the lack of correlation comes from one main cause. Discriminating transcripts of a same gene that share common sequences with short reads is difficult. Longer reads are clearly helpful, however they do not always enable to discriminate transcripts. Indeed, in the case where a read only covers the 3’ end of a transcript, and not the full length, it may be ambiguously assigned to several transcripts.

For example, for the *Swi5* gene, although several rare (lowly expressed) transcripts are seen only with Illumina, the other ones are harder to quantify (red dots in figure 5a). RSEM uses the unique part of each transcript to proportionally allocate the reads that mapped equally on the common part of the transcript. In the case where a transcript has no read which uniquely maps to it, its expression cannot be computed and is set to 0. This is the case for the transcript ENSMUST00000050410 (*Swi5*-201, Supplementary figure 9) of *Swi5*, whose expression is underestimated (0 TPM). Conversely, some transcripts are underestimated by ONT RNA-Seq. This is the case of *Swi5*-204, whose unique region is located at the 5’ end of the gene, and is therefore poorly covered by long reads.

To avoid the difficult step of correctly assigning a read to a transcript, we summed the quantification of all transcripts for each gene. Figure 5c shows the quantification at gene level. As reported in other papers [8] [5] [24] the correlation at gene level is quite good (*ρ* =0.78). However inter-genes repeats remains a cause of mis-quantified genes. For example, a large part of the mitochondrial chromosome had been recently integrated in the mouse chromosome 1 [15]. As a consequence, 7 genes are present in 2 copies in the genome, one copy annotated as functional on the mitochondrial chromosome and another one, annotated as pseudogene on chromosome 1 (shown in red in figure 5c). Since this integration is recent, the copies did not diverge yet. They are therefore difficult to quantify due to multimapping, even when using long reads, since the repeat is larger than the full transcript.

### Quantification of processed pseudogenes

These are particular cases of processed pseudogenes which come from the retrotranscription and reintegration in the genome of one of the transcript of their parent gene[11]. After their integration, they have no intron and, without any selective pressure, they diverge from their parent gene proportionally with their age. Some of them are expressed [6] and are annotated as transcribed processed pseudogenes although the vast majority of pseudogenes are not expressed [11]. Correctly assigning the reads to the parent gene and not the pseudogene is not trivial.

Figure 5d shows that mapping long reads to the genome with Minimap 2 (-ax splice) (as used in [30]) results in the mis-quantification of processed pseudogenes (green points in Figure 5d). The expression of most of them is over-estimated by the ONT RNA-Seq protocol (this is also the case of ONT cDNA-Seq). It can be explained by two main reasons.

First, it can come from the fact that if a mapper has to choose between two genomic locations, one with gaps (introns of the parent gene), and one with no gaps (the processed pseudogene), it will tend to favour the gapless mapping, as gapless alignment are easier to find. We note that the scoring system of minimap2 consists in selecting the max-scoring sub-segment, excluding introns, and therefore not explicitly favouring the gapless mapping. However, this requires that splice sites are correctly identified in the first place, a task which remains difficult with noisy long reads.

A second reason explaining the overestimation of processed pseudogenes is related to polyA tails. Processed pseudogenes originate from transcripts which contained a polyA tail, which was then integrated in the genome, downstream the pseudogene. Many of the ONT reads originating from the parent gene also contain this polyA tail, favoring the alignment at the processed pseudogene genomic location. The alignment will be longer thanks to the polyA tail. An example is shown in figure 6. The processed pseudogene *Rpl17-ps8* differs from its parent gene by two bases (A to G at the position chrX:96,485,078 and A to G at the position chrX:96,485,267). These divergences are marked in red in the figure. At these two positions we observe that reads differ from the reference genome : they have a G instead of an A. This means that these reads come from the parent gene and we mistakenly aligned them onto the pseudogene because it is intronless and contains a polyA tail.

**Figure 6:**
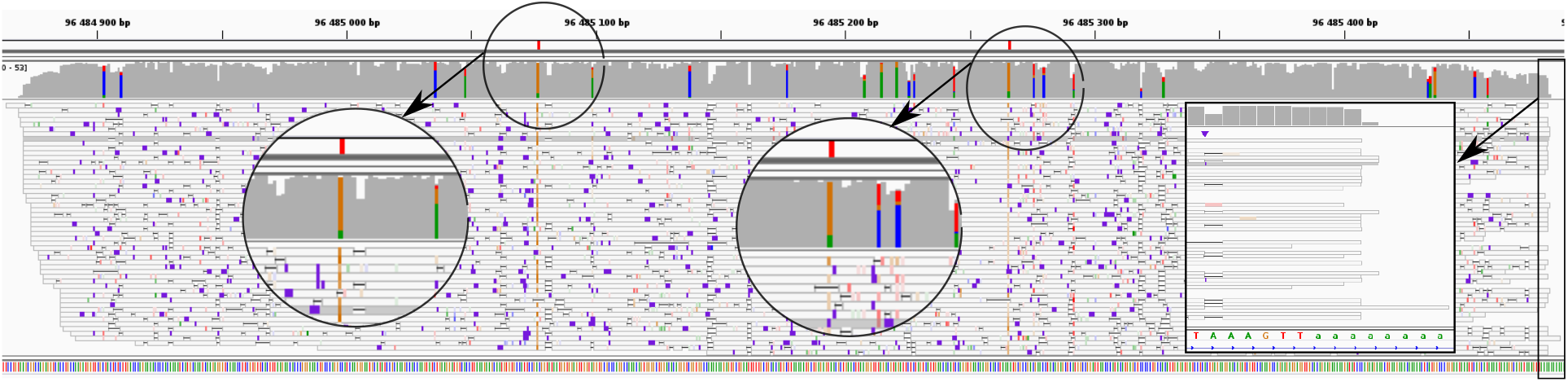
Example of a processed pseudogene whose expression is overestimated : *Rpl17-ps8* (retro Rpl17) Alignment visualization with IGV of *Rpl17-ps8*. The positions of divergence between Rpl17-ps8 and Rpl17 are shown in red in the first track. Second track is ONT RNA-Seq coverage, third track is ONT RNA-Seq reads. Colored positions in the coverage track correspond to mismatches. Most reads contain mismatches at the exact position of the divergences with the parent gene. They are therefore incorrectly mapped, partly because they overlap the polyA tail which is integrated in the genome downstream the pseudogene.

### Quantification of genes overlapping transposable elements

Another example of repeat-associated gene biotype is given by the long non-coding RNAs (lncRNAs) which are highly enriched in transposable elements (TEs). These TEs are sometimes considered as the functional domains of lncRNAs [10], and it has been estimated that 66% of mouse lncRNA overlap at least one TE [12]. Our experimental design allows us to assess the impact of read lengths on noncoding (versus protein-coding) gene quantifications for different levels of TE coverage. As expected, the higher the TE content of a gene, the larger the difference in quantification between long and short-read sequencing technologies (Supplementary Figure 10a). Although this tendency is observed for both protein-coding and lncRNAs biotypes, lncRNAs are more impacted given that they are more prone to be enriched in TEs. One interesting example is given by the known imprinted lncRNA *KCNQ1OT1* (ENSMUSG00000101609) which is specifically expressed from the paternal allele in opposite direction to the *KCNQ1* protein-coding gene [20] (Supplementary Figure 10b). About 41% of the *KCNQ1OT1* transcript sequence is composed of TE elements and its quantification using Illumina TruSeq versus ONT cDNA-Seq protocols highlights contrasting values (TPM = 0.36 and ONT cDNA-Seq = 118).

## Discussion

In this work, we generated a dataset which we think should be of general interest for the community. This dataset consists of RNA and cDNA sequencing of the same samples using both Illumina and ONT technologies. Importantly, we also sequenced Lexogen E2 spike-in data, together with our mouse samples, which enabled us to assess which technology yielded the most accurate quantification. Although lexogen spike-in have been used to evaluate the quantification obtained with ONT cDNA-Seq [29] or ONT RNA-Seq [8] protocol separately, we are the first to compare the quantification obtained with ONT cDNA-Seq, RNA-Seq and Illumina cDNA-Seq.

Using the spike-in data, we find that the ONT RNA-Seq protocol is the most accurate, slightly better than the widely used Illumina TruSeq protocol. In contrast, the cDNA-Seq data was more biased and yielded a poorer quantification.

We further found that transcripts with internal runs of poly(T) tend to be truncated and oversampled when using the ONT cDNA-Seq protocol. Sequencing the same library preparation with the Illumina technology enabled us to confirm that the truncation issue was related to the sample preparation and not to the sequencing. We further show that this bias is not restricted to our dataset, and can be found in a human ONT dataset [30]. Truncation biases associated to internal runs of poly(A) had been reported earlier and motivated the usage of anchored poly-dT primers (poly-TVN) [21]. On the other hand, biases associated to internal runs of poly(T) had remained undetected, although they may affect more than 20% of expressed transcripts in mouse. This bias could also affect other long-reads cDNA-Seq data. Although biases had been searched for in previous work [13], it may have remained undetected because the authors were then focusing on internal runs of at least 20 A’s.

We then used our data to quantify mouse genes and found that ONT RNA-Seq quantification correlated well with Illumina cDNA-Seq quantification (Figure 5c) but when trying to quantify at the transcript level, the correlation was overall poorer (Figure 5a). A temptation could be to think that ONT RNA-Seq yields better transcript-level quantification as reads are longer and are, unlike short reads, unambiguously assigned to a single transcript. In practice, 70% of ONT RNA-Seq reads are assigned to a single transcript, while the remaining 30% are ambiguously mapped. This was particularly the case for transcripts which differed at their 5’end, like in Swi5. Quantifying transcripts and not genes is still challenging, and requires the development of dedicated bioinformatics methods. When trying to use salmon [22] on long reads, as in [25], we did not obtain better results than when simply counting primary alignments. There should however be room for improvement in this direction, and our spike-in dataset could be a good training set for future methods.

In this work, we chose to align reads to a reference transcriptome. Indeed, when trying to map reads to the reference genome, we observed a systematic over-estimation of the quantification of processed pseudogenes, at the expense of their parent gene. We further show that this biased quantification is due to alignment issues: 1-poly(A) tails of pseudogenes are integrated in the genome and ‘attract’ reads from the parent gene and 2-accurate identification of splice sites when mapping long RNA-Seq reads is challenging, which disfavors the parent gene.

We therefore strongly recommend to map reads on the reference transcriptome and not on the genome, as reference transcripts do not contain introns, nor poly(A) tails. However, a clear limitation of aligning reads to a reference annotation, instead of a reference genome, is that we cannot discover novel transcripts. As a consequence, reads stemming from these novel transcripts will be unmapped, or incorrectly assigned to alternative transcripts (as in APOE gene, Supplementary Figure 11). Improving alignment tools to correctly handle processed pseudogenes seems essential to identify and quantify transcripts, especially in the case of non-model species where no exhaustive annotation is available. More generally, the quantification of repeat-containing genes is difficult. Long reads are particularly useful for quantifying these genes, like long non coding RNAs, which are enriched in transposable elements.

There is currently a lot of interest for the potential of ONT RNA-Seq to identify and quantify genes and transcripts, as can be seen by the currently low but expanding number of datasets available with this technology. Here we proposed the first dataset on mouse with several interesting and unique features, as Lexogen E2 spike-ins, Illumina sequencing of ONT library preparation or Lexogen TeloPrime protocol. We think that ONT sequencing is promising for studying RNA, especially if the number of reads and full-length reads continues to increase. Improvements in the technology and library preparation protocol to obtain more reads and more full-length reads are also expected to be very helpful in obtaining precise quantification of all genes and transcripts. The recent launch of the PromethION device will allow a deep sequencing of transcriptomes which should enable to overcome the limitations of the MinION device.

## Methods

### Biological material

We used total RNA extracted from mouse brain (Cat # 636601, lot number 1403636A and 1605262A) and liver (Cat # 636603, lot number 1305118A) from Clontech (Mountain View, CA, USA).

### Libraries preparation

#### Illumina cDNA library

RNA-Seq library preparations were carried out from a mix of 250 ng total RNA and 0.25 ng Spike-in RNA Variant Control Mix E2 (Lexogen, Vienna, Austria) using the TruSeq Stranded mRNA kit (Illumina, San Diego, CA, USA), which allows mRNA strand orientation. Ready-to-sequence Illumina libraries were quantified by qPCR using the KAPA Library Quantification Kit for Illumina Libraries (KapaBiosystems, Wilmington, MA, USA), and libraries profiles evaluated with an Agilent 2100 Bioanalyzer (Agilent Technologies, Santa Clara, CA, USA).

#### Illumina on Nanopore cDNA library

250 ng of cDNA prepared using the “cDNA-PCR Sequencing” protocol (see “Nanopore cDNA library” below) were sonicated to a 100-to 1000-bp size using the E220 Covaris instrument (Covaris, Woburn, MA, USA). Fragments were end-repaired, then 3’-adenylated, and NEXTflex PCR free barcodes adapters (Bioo Scientific, Austin, TX, USA) were added using NEBNext Sample Reagent Module (New England Biolabs, Ipswich, MA, USA). Ligation products were amplified using Illumina adapter-specific primers and KAPA HiFi Library Amplification Kit (KapaBiosystems, Wilmington, MA, USA) and then purified with AMPure XP beads (Beckmann Coulter, Brea, CA, USA). Ready-to-sequence Illumina libraries were quantified by qPCR using the KAPA Library Quantification Kit for Illumina Libraries (KapaBiosystems), and libraries profiles evaluated with an Agilent 2100 Bioanalyzer (Agilent Technologies).

#### Nanopore cDNA library

Total RNA was first depleted using the Ribo-Zero rRNA Removal Kit (Human/Mouse/Rat) (Illumina). RNA was then purified and concentrated on a RNA Clean Concentrator^TM^-5 column (Zymo Research, Irvine, CA, USA). cDNA libraries were performed from a mix of 50ng RNA and 0.5 ng Spike-in RNA Variant Control Mix E2 (Lexogen) according to the Oxford Nanopore Technologies (Oxford Nanopore Technologies Ltd, Oxford, UK) protocol “cDNA-PCR Sequencing” with a 14 cycles PCR (8 minutes for elongation time). ONT adapters were ligated to 650 ng of cDNA.

#### Nanopore RNA library

RNA libraries were performed from a mix of 500ng RNA and 5ng Spike-in RNA Variant Control Mix E2 (Lexogen) according to the ONT protocol “Direct RNA sequencing”. We performed the optional reverse transcription step to improve throughput, but cDNA strand was not sequenced.

#### Nanopore TeloPrime library

Three cDNA libraries were performed from 2*μ*g total RNA for each RNA sample according to the TeloPrime Full-Length cDNA Amplification protocol (Lexogen). A total of 5 PCR were carried out with 30 to 40 cycles for the brain sample and 30 cycles for the liver sample. Amplifications were then pooled and quantified. Nanopore libraries were performed from respectively 560ng and 1000ng of cDNA using the SQK-LSK108 kit according to the Oxford Nanopore protocol.

### Sequencing and reads processing

#### Illumina datasets

Illumina cDNA libraries, prepared with the TruSeq (TruSeq_SR) and Nanopore (PCS108_SR) protocols, were sequenced using 151 bp paired end reads chemistry on a HiSeq4000 Illumina sequencer (Table 1). After the Illumina sequencing, an in-house quality control process was applied to the reads that passed the Illumina quality filters. The first step discards low-quality nucleotides (Q < 20) from both ends of the reads. Next, Illumina sequencing adapters and primer sequences were removed from the reads. Then, reads shorter than 30 nucleotides after trimming were discarded. The last step identifies and discards read pairs that mapped to the phage phiX genome, using SOAP [18] and the phiX reference sequence (GenBank: NC_001422.1). These trimming and removal steps were achieved using in-house-designed software as described in [2].

#### Nanopore datasets

Nanopore libraries were sequenced using a MinION Mk1b with R9.4.1 (PCS108_LR and RNA001_LR) or R9.5 flowcells (TELO_LR). The data were generated using MinKNOW 1.11.5 and basecalled with Albacore 2.1.10 (Table 1).

### Reads alignment and transcripts quantification

Long reads were mapped to the spike-in transcripts using Minimap2 (version 2.14) [17] (-ax map-ont). Supplementary alignments, secondary alignments and reads aligned on less than 80% of their length were filtered out. We used the number of aligned reads as a proxy of the expression of a given transcript. Short reads were mapped to the spike-in transcripts using bowtie [14] and quantified using RSEM [16]. The quantification obtained is given in TPM (transcript per million).

We then assessed the mouse transcripts expression and mapped the long reads against the mouse transcripts (Ensembl 94) using Minimap2 (with the following options -ax map-ont and -uf for direct RNA reads). Long reads from cDNA (PCS108_LR) and TeloPrime (TELO_LR) were trimmed using porechop and default parameters before alignment against the mouse transcripts. Long reads from RNA (RNA001_LR) were not trimmed, as the ONT basecaller could not detect DNA adapters. Supplementary alignements, secondary alignments and reads aligned on less than 80% of their length were filtered out. Expression was directly approximated by the number of reads which mapped on a given transcript. Long reads were also mapped on the reference genome using Minimap2 (-ax splice). Supplementary alignments, secondary alignments and reads aligned on less than 80% of their length were filtered out Short reads were mapped to the reference genome (release Grcm38.p6) using STAR with the gtf option (annotation Ensembl 94). In order to quantify each transcript, short reads were also mapped on the referencence transcriptome using bowtie and quantification were obtained with RSEM.

### Evaluating the ability of each technology to predict the true SIRV quantification using cross-validation

We build 3 models: *M*_1_ : *log*(*SIRV*) = *μ*_1_ + *β*_1_ * *log*(*readCountcDNA*) + *error*; *M*_2_ : *log*(*SIRV*) = *μ*_2_ + *β*_2_ * *log*(*TPM*) + *error*; *M*_3_ : *log*(*SIRV*) = *μ*_3_ + *β*_3_ * *log*(*readCountRNA*) + *error*. As these models are not nested, they cannot be compared against each other with likelihood ratio tests. We therefore use cross-validation, using 4/5 of our 67 SIRV to estimate the parameters of each model, and the remaining 1/5 to estimate the quality of the prediction. We repeat this process 30 times, choosing randomly a different partition to train and test the model, and we obtain confidence intervals on the prediction error for each model.

#### Truncated Reads Analysis

For each transcript annotated in Ensembl94 containing an internal run of at least 9Ts, we computed the number of reads covering the following positions: 25, 50, 75 and 100nt upstream and downstream the internal run of poly(T). For each transcript *t*, the most covered position was retrieved, and the number of reads covering this position was noted *max_t_*. The coverage of each position was then divided by *max_t_*, so as to obtain a normalised coverage. Then for each position, we computed the mean of the relative coverage at this position across all transcripts verifying *max_t_* > 10. This is the value plotted in Figure 3a. The error bars represent the standard error around the mean. The same analysis was done for the human ONT dataset, using gencode27 annotations (Supplementary Figure 2).

#### Saturation curve

For short reads, we kept only the best alignment as reported by RSEM and the primary alignment of each long read. Only protein coding transcripts (transcript_biotype=protein_coding) were taken into account.

### Quantification of TE-containing genes

Given that lncRNAs are lowly expressed, for this specific analysis, we restricted to cDNA-Seq and did not apply our 80% query coverage filter. Using annotated TEs from the RepeatMasker database [26], we classified lncRNAs and mRNAs based on their TE coverage in four categories (with the “0%” class corresponding to genes without any exonic-overlapping TE and conversely, the class of “>66-100%” for genes highly enriched in exonic TE). For each expressed gene, we further computed the ratio between Nanopore cDNA versus Illumina TruSeq gene quantifications with respect to their TE categories.

## Availability of supporting data

The Illumina and MinION data are available in the European Nucleotide under the following accession number PRJEB27590. The entire dataset (fastq and bam files) is available from the following website: http://www.genoscope.cns.fr/externe/ONT_mouse_RNA.

## Acknowledgements

The authors are grateful to Oxford Nanopore Technologies Ltd for providing early access to the MinION device through the MAP, and we thank the staff of Oxford Nanopore Technology Ltd for technical help, particularly Botond Sipos, Daniel Turner, Michelle Hiscutt and James Platt for insightful discussions about poly-T truncation phenomenon. We thank Philippe Veber and Arnaud Mary for their advice on statistical and bioinformatics analyses.

## Authors contribution

CC optimized and performed the sequencing. CS, CDS, AC, MD, TD, VL and JMA performed the bioinformatic analyses. CS, VL and JMA wrote the article. VL and JMA supervised the study.

## References

1. Alberti, A. et al. Comparison of library preparation methods reveals their impact on interpretation of metatranscriptomic data. BMC Genomics 15, 912. ISSN: 1471-2164 (Oct. 2014).

2. Alberti, A. et al. Viral to metazoan marine plankton nucleotide sequences from the Tara Oceans expedition. Scientific Data 4, 170093. ISSN: 2052-4463 (Aug. 2017).

3. Belser, C. et al. Chromosome-scale assemblies of plant genomes using nanopore long reads and optical maps. Nature Plants 4, 879–887. ISSN: 2055-0278 (Nov. 2018).

4. Brooks, A. (Nanopore RNA Consortium) - Native RNA sequencing of human polyadenylated transcripts https://nanoporetech.com/resource-centre/native-rna-sequencing-human-polyadenylated-transcripts. [Accessed 25 Fev 2019]. 2018.

5. Byrne, A. et al. ARTICLE Nanopore long-read RNAseq reveals widespread transcriptional variation among the surface receptors of individual B cells. Nature Communications 8. doi:10.1038/ncomms16027. https://www.nature.com/articles/ncomms16027.pdf (2017).

6. Carelli, F. N. et al. The life history of retrocopies illuminates the evolution of new mammalian genes. Genome research 26, 301–14. ISSN: 1549-5469 (Mar. 2016).

7. Deamer, D., Akeson, M. & Branton, D. Three decades of nanopore sequencing. Nature Biotechnology 34, 518–524. ISSN: 1087-0156 (May 2016).

8. Garalde, D. R. et al. Highly parallel direct RNA sequencing on an array of nanopores. Nature Methods 15, 201–206. ISSN: 1548-7091 (Jan. 2018).

9. Jain, M. et al. Nanopore sequencing and assembly of a human genome with ultra-long reads. Nature Biotechnology 36, 338–345. ISSN: 1087-0156 (Jan. 2018).

10. Johnson, R. & Guigo, R. The RIDL hypothesis: transposable elements as functional domains of long noncoding RNAs. RNA 20, 959–976. ISSN: 1355-8382 (July 2014).

11. Kaessmann, H., Vinckenbosch, N. & Long, M. RNA-based gene duplication: mechanistic and evolutionary insights. Nature reviews. Genetics 10, 19–31. ISSN: 1471-0064 (Jan. 2009).

12. Kelley, D. & Rinn, J. Transposable elements reveal a stem cell-specific class of long noncoding RNAs. Genome Biology 13, R107. ISSN: 1465-6906 (2012).

13. Kuo, R. I. et al. Normalized long read RNA sequencing in chicken reveals transcriptome complexity similar to human. BMC Genomics 18, 323. ISSN: 1471-2164 (Apr. 2017).

14. Langmead, B. Aligning short sequencing reads with Bowtie. Current protocols in bioinformatics Chapter 11, Unit 11.7. ISSN: 1934-340X (Dec. 2010).

15. Leister, D. & Richly, E. NUMTs in Sequenced Eukaryotic Genomes. Molecular Biology and Evolution 21, 1081–1084. ISSN: 0737-4038 (June 2004).

16. Li, B. & Dewey, C. N. RSEM: accurate transcript quantification from RNA-Seq data with or without a reference genome. BMC Bioinformatics 12, 323. ISSN: 1471-2105 (Dec. 2011).

17. Li, H. Minimap2: pairwise alignment for nucleotide sequences. Bioinformatics 34, 3094–3100 (2018).

18. Li, R. et al. SOAP2: an improved ultrafast tool for short read alignment. Bioinformatics 25, 1966–1967. ISSN: 1367-4803 (Aug. 2009).

19. Lipson, D. et al. Quantification of the yeast transcriptome by single-molecule sequencing. Nature Biotechnology 27, 652–658. ISSN: 1087-0156 (July 2009).

20. Mancini-DiNardo, D., Steele, S. J. S., Levorse, J. M., Ingram, R. S. & Tilghman, S. M. Elongation of the Kcnq1ot1 transcript is required for genomic imprinting of neighboring genes. Genes & Development 20, 1268–1282. ISSN: 0890-9369 (May 2006).

21. Nam, D. K. et al. Oligo(dT) primer generates a high frequency of truncated cDNAs through internal poly(A) priming during reverse transcription. Proceedings of the National Academy of Sciences of the United States of America 99, 6152–6. ISSN: 0027-8424 (Apr. 2002).

22. Patro, R., Duggal, G., Love, M. I., Irizarry, R. A. & Kingsford, C. Salmon provides fast and bias-aware quantification of transcript expression. Nat. Methods 14, 417–419 (Apr. 2017).

23. Schmidt, M. H.-W. et al. De Novo Assembly of a New Solanum pennellii Accession Using Nanopore Sequencing. The Plant cell 29, 2336–2348. ISSN: 1532-298X (Oct. 2017).

24. Seki, M. et al. Evaluation and application of RNA-Seq by MinION. DNA Research, dsy038 (2018).

25. Soneson, C. et al. A comprehensive examination of Nanopore native RNA sequencing for characterization of complex transcriptomes. bioRxiv. doi:10.1101/574525. eprint: https://www.biorxiv.org/content/early/2019/03/11/574525.full.pdf. https://www.biorxiv.org/content/early/2019/03/11/574525 (2019).

26. Tarailo-Graovac, M. & Chen, N. Using RepeatMasker to identify repetitive elements in genomic sequences. Current protocols in bioinformatics Chapter 4, Unit 4.10. ISSN: 1934-340X (Mar. 2009).

27. Van Dijk, E. L., Jaszczyszyn, Y. & Thermes, C. Library preparation methods for next-generation sequencing: Tone down the bias. Experimental Cell Research 322, 12–20. ISSN: 00144827 (Mar. 2014).

28. Wang, Z., Gerstein, M. & Snyder, M. RNA-Seq: a revolutionary tool for transcriptomics. Nature reviews. Genetics 10, 57–63. ISSN: 1471-0064 (Jan. 2009).

29. Weirather, J. L. et al. Comprehensive comparison of Pacific Biosciences and Oxford Nanopore Technologies and their applications to transcriptome analysis. F1000Research 6, 100. ISSN: 20461402 (Feb. 2017).

30. Workman, R. E. et al. Nanopore native RNA sequencing of a human poly(A) transcriptome. bioRxiv. doi:10.1101/459529. eprint: https://www.biorxiv.org/content/early/2018/11/09/459529.full.pdf. https://www.biorxiv.org/content/early/2018/11/09/459529 (2018).

