## Supplementary Information for "Transcriptome profiling of mouse samples using nanopore sequencing of cDNA and RNA molecules"

### Supplementary Figures

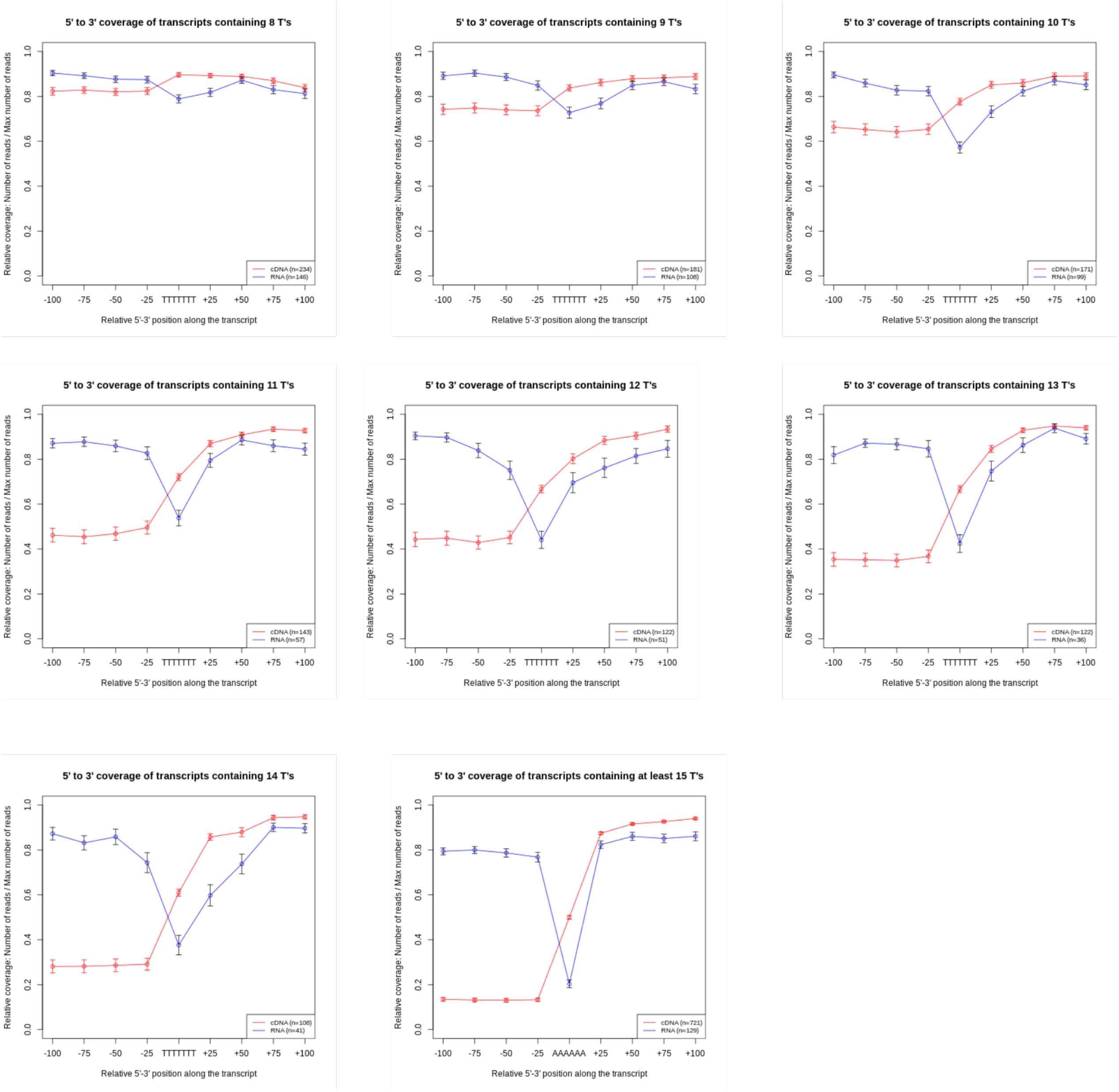


Figure 1: **Poly(T) induce 5' truncated reads** Relative coverage of transcripts for our ONT cDNA-Seq dataset and our ONT RNA-Seq dataset for transcripts covered by at least 10 reads around a poly(T). Several size of poly(T) have been tested and we found that the effect is visible using the cDNA-Seq dataset from poly(T) longer than 9 T's : transcripts containing stretches of at least 9 T's are less covered in 5' than other transcripts. In all cases, the local coverage deficit observed in the ONT RNA-seq dataset is due to sequencing error causing by the homopolymers.

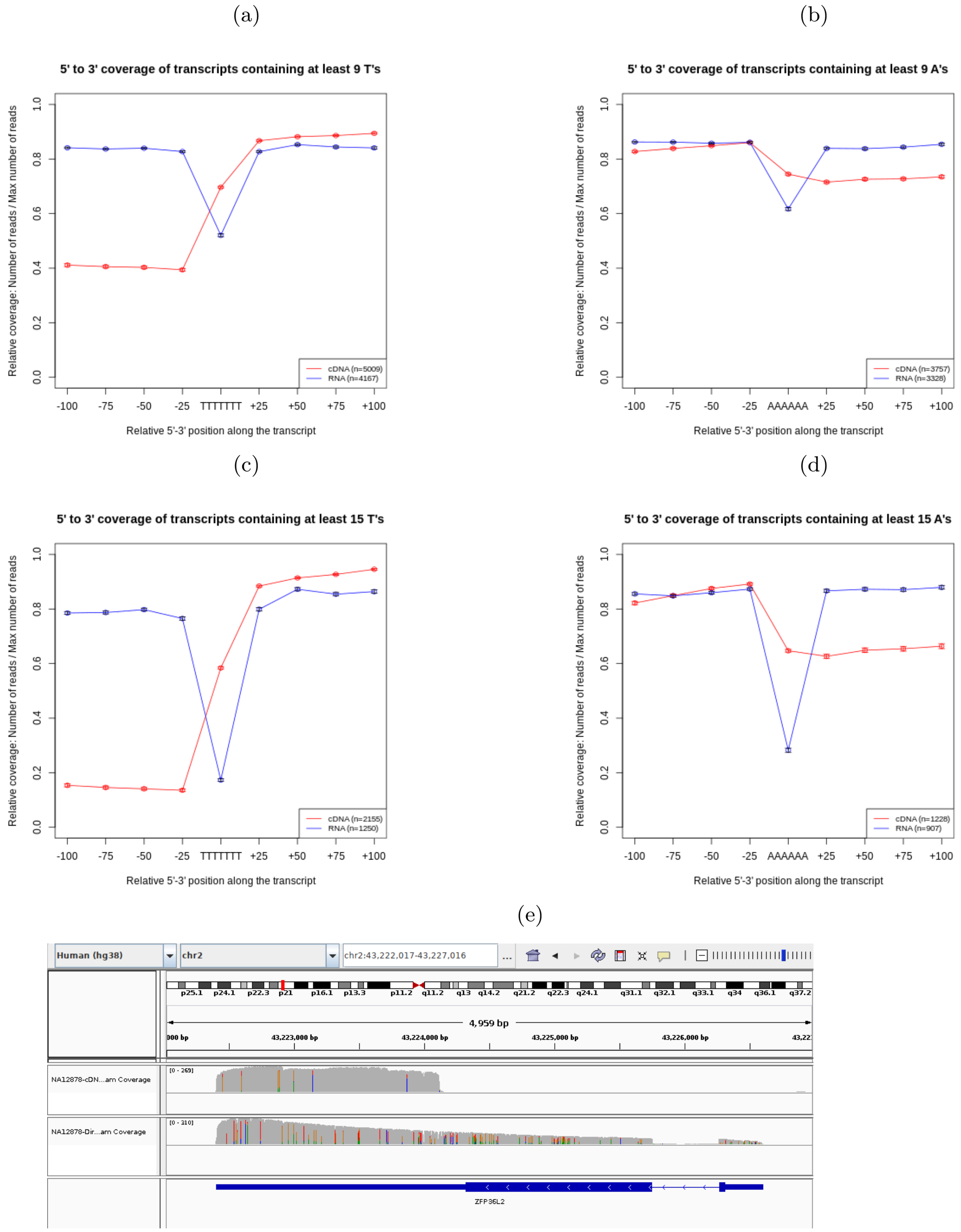


Figure 2: **Truncated Reads** Relative coverage of transcripts for the ONT cDNA-Seq dataset and the ONT RNA-Seq dataset of Workman et al. for transcripts covered by at least 10 reads around an internal run of poly(T) (panels a and c) or poly(A) (panels b and d). Using the ONT CDNA-Seq dataset, transcripts containing internal runs of poly(T) are less covered in 5' than other transcripts, wheras transcripts containing internal runs of poly(A) are less covered in 3'. It indicates that these transcripts are covered by a high proportion of truncated reads. The coverage deficit observed in the ONT RNA-seq dataset is due to sequencing errors caused by the homopolymers. The effect is more marked when considering internal runs of at least 15 T's (panel c) or 15 A's (panel d). (e) Example obtained using the Workman et al. dataset. The *ZFP36L2* gene contains an internal run of 11 T's. Reads from the ONT cDNA-Seq are truncated (first track) whereas ONT RNA-Seq reads are not (second track).

(a)

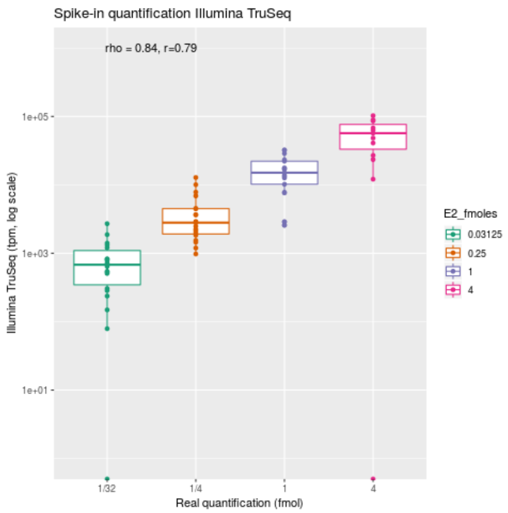


(b)

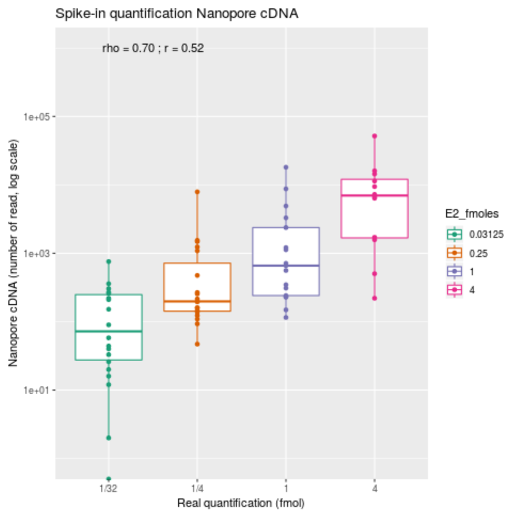


(c)

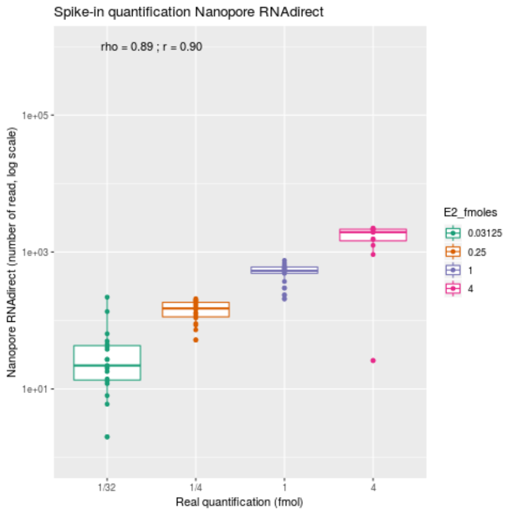


Figure 3: **Spike in quantification for C2R1 datasets** Reads were mapped against the SIRV transcriptome and quantifications computed at transcript level. The observed quantification are correlated with the known theoretical quantification of the spike in. (a) Correlation obtained for Illumina with the TruSeq protocol (Spearman's rho =0.84 ). (b) Corelation obtained for Nanopore with the cDNA protocol (Spearman's rho = 0.70 ). (c) Corelation obtained for Nanopore with the RNA direct protocol (Spearman's rho = 0.89 ).

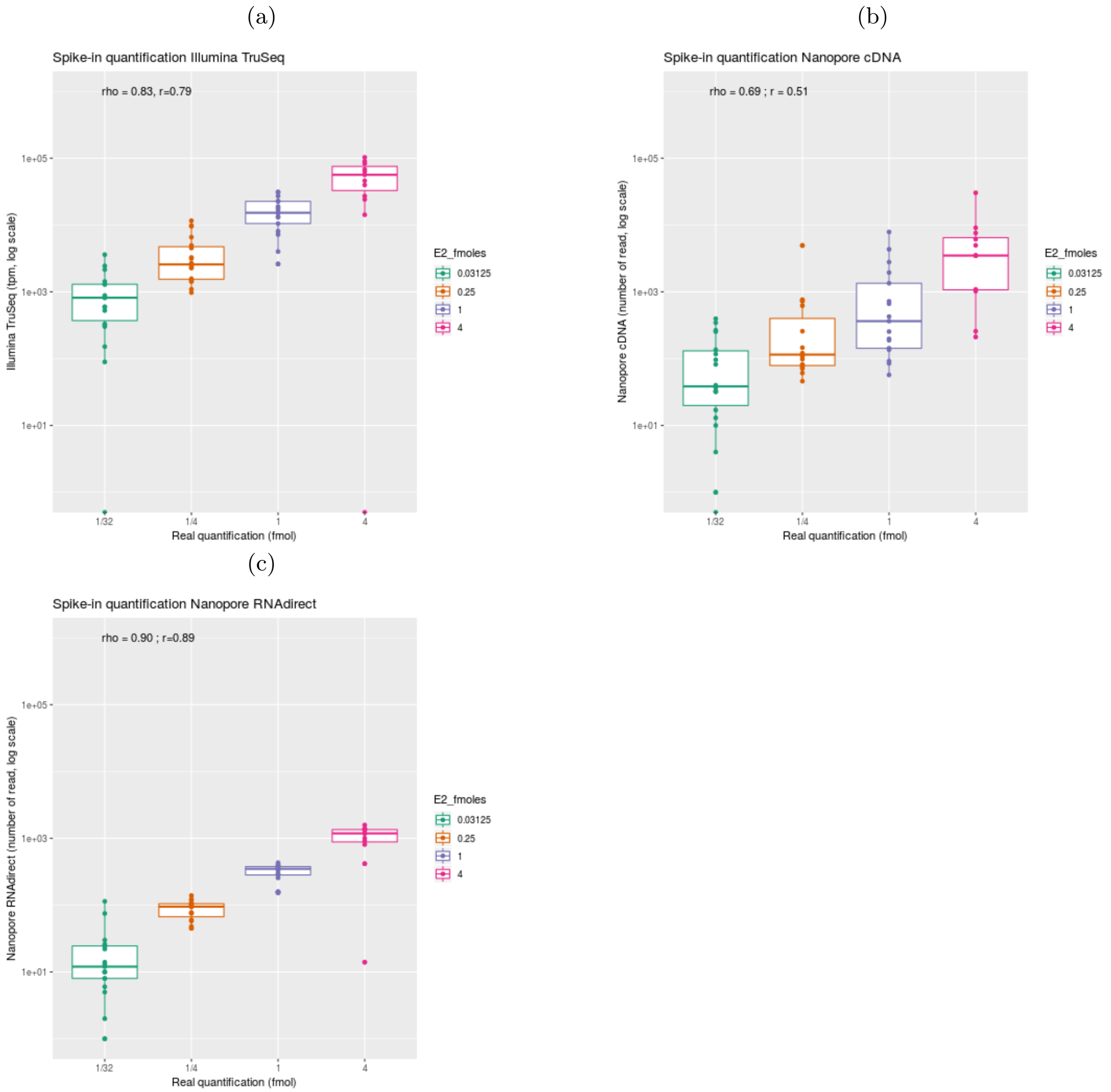


Figure 4: **Spike in quantification for C2R2 datasets** Reads were mapped against the SIRV transcriptome and quantifications computed at transcript level. The observed quantification are correlated with the known theoretical quantification of the spike in. (a) Correlation obtained for Illumina with the TruSeq protocol (Spearman's rho = 0.83). (b) Corelation obtained for Nanopore with the cDNA protocol (Spearman's rho = 0.69 ). (c) Corelation obtained for Nanopore with the RNA direct protocol (Spearman's rho = 0.90 ).

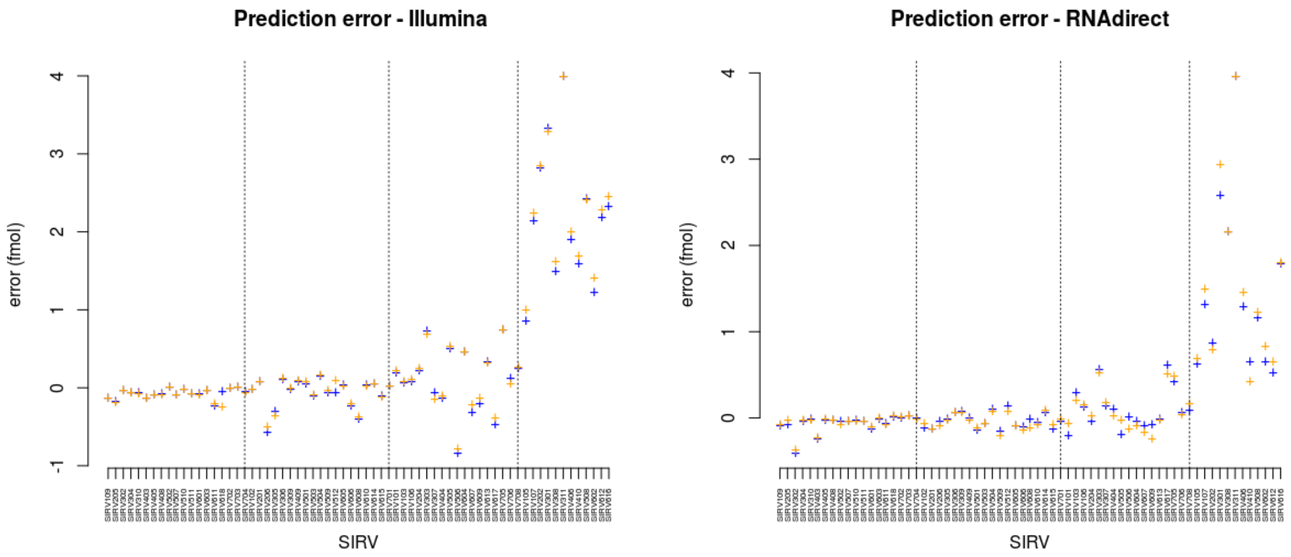


Figure 5: **Reproductibility of the prediction error.** The error between the prediction and the real quantification has bee computed for each replicates C2R1 and C2R2 for (a) the illumina dataset and (b) the Nanopore RNAdirect dataset.

(a)

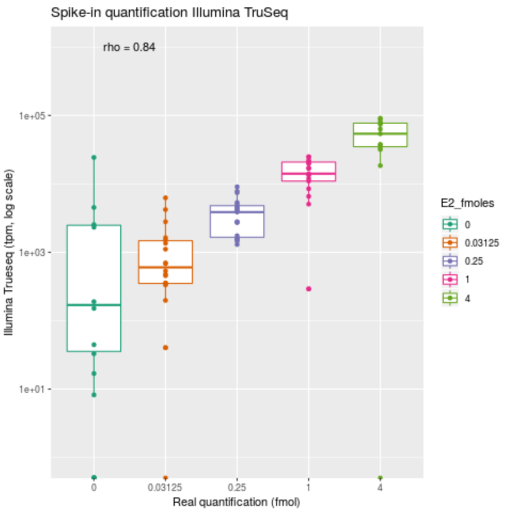


(b)

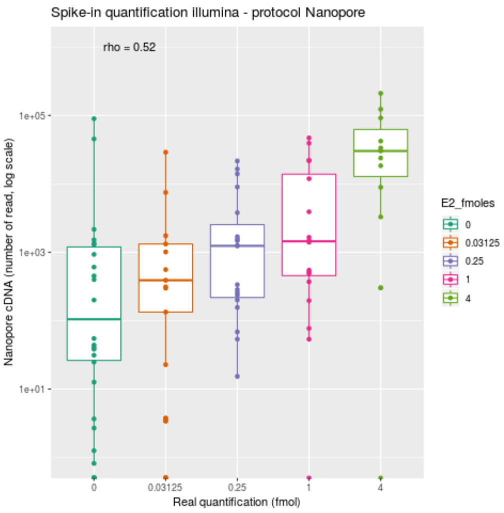


(c)

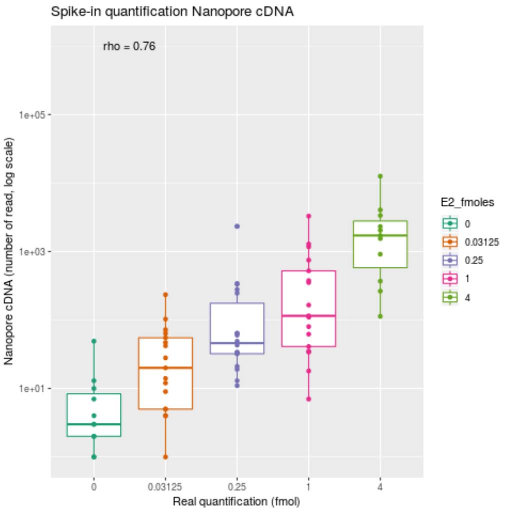


(d)

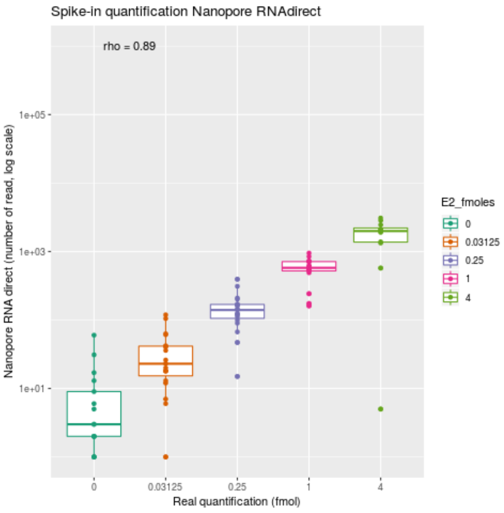


Figure 6: **Evaluation of quantification using the SIRV E2 spike-in mix and the over-annotation supplied by Lexogen** Reads were mapped against the SIRV transcriptome and quantifications computed at transcript level. The observed quantification are correlated with the known theoretical quantification of the spike in. (a) Correlation obtained for Illumina with the TruSeq protocol (Spearman's rho = 0.84). (b) Correlation obtained for illumina with the cDNA synthesis Nanopore protocol (Spearman's rho =0.52). (c) Corelation obtained for Nanopore with the cDNA protocol (Spearman's rho = 0.76). (d) Corelation obtained for Nanopore with the RNA direct protocol (Spearman's rho = 0.89).

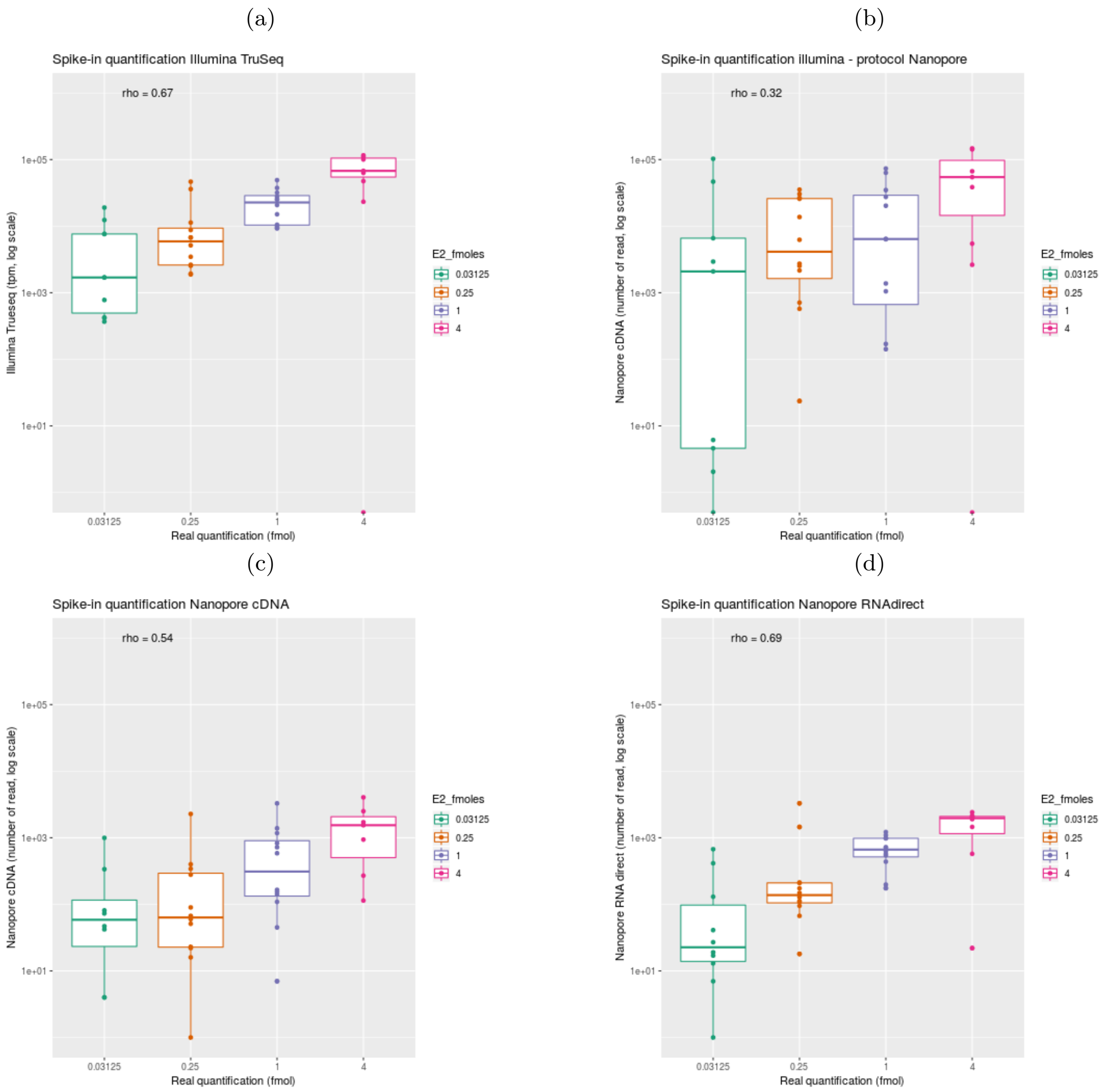


Figure 7: **Evaluation of quantification using the SIRV E2 spike-in mix and the incomplete annotation supplied by Lexogen** Reads were mapped against the SIRV transcriptome and quantifications computed at transcript level. The observed quantification are correlated with the known theoretical quantification of the spike in. (a) Correlation obtained for Illumina with the TruSeq protocol (Spearman's rho = 0.67). (b) Correlation obtained for illumina with the cDNA synthesis Nanopore protocol (Spearman's rho =0.32). (c) Corelation obtained for Nanopore with the cDNA protocol (Spearman's rho = 0.54). (d) Corelation obtained for Nanopore with the RNA direct protocol (Spearman's rho = 0.69).

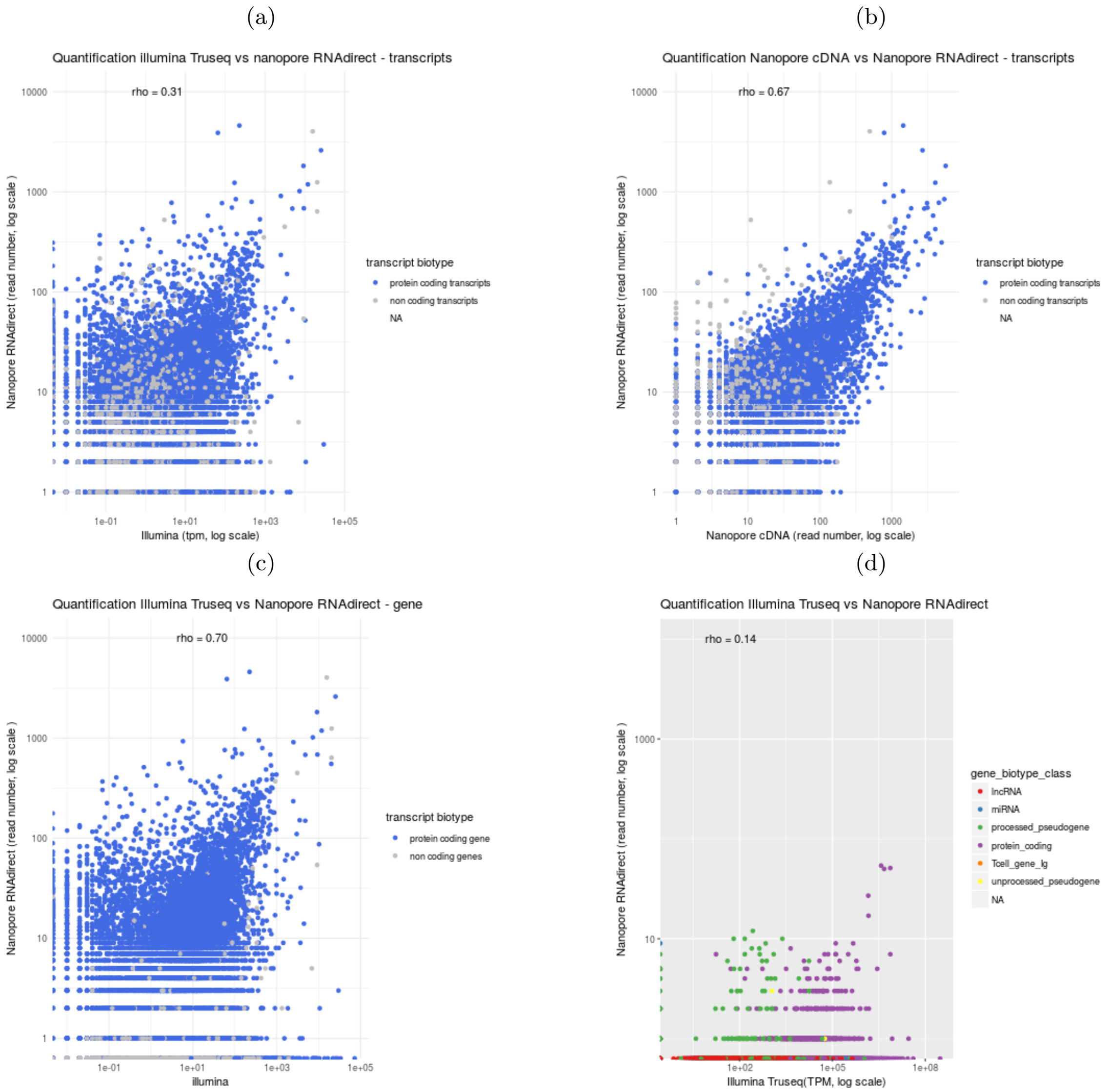


Figure 8: **Comparison of quantifications in Liver** Reads were mapped against the mouse reference transcriptome. Transcript annotated as coding protein transcript are in blue. Spearman's $\rho$ has been computed for all transcripts. (a) Comparison of Nanopore RNA direct and Illumina (TruSeq) quantifications (Spearman's $\rho = 0.31$). (b) Comparison of Nanopore RNA direct and Nanopore cDNA quantifications (Spearman's $\rho = 0.67$). (c) Comparison of Nanopore RNA direct and Illumina (TruSeq) quantifications. Transcript quantification were summed for each gene. (Spearman's $\rho = 0.70$). (d) Reads were mapped against the mouse reference genome and quantifications computed at gene level. We compared the Nanopore RNAdirect and the illumina Truseq protocols (Spearman's $\rho$ $= 0.14$). Green points correspond to processed pseudogenes.

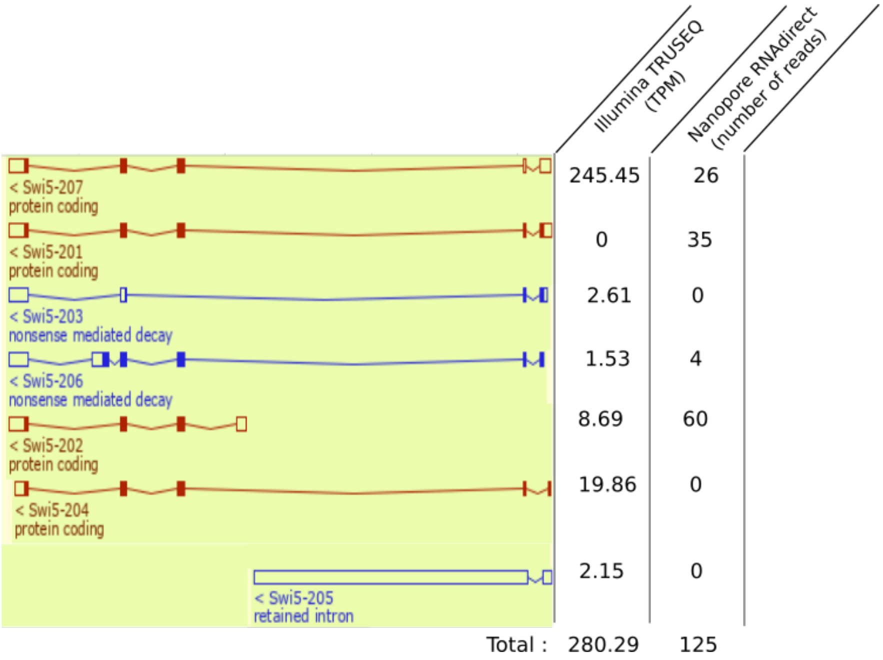


Figure 9: **Quantifications for Swi5 trancripts.** Swi5 annotation visualized with the Ensembl genome browser. The transcript Swi5-201 has no short read which uniquely maps to it. Therefore RSEM cannot allocate read to this transcript. With ONT RNA-Seq we have long enough reads to distinguish it from the other transcripts.

(a)

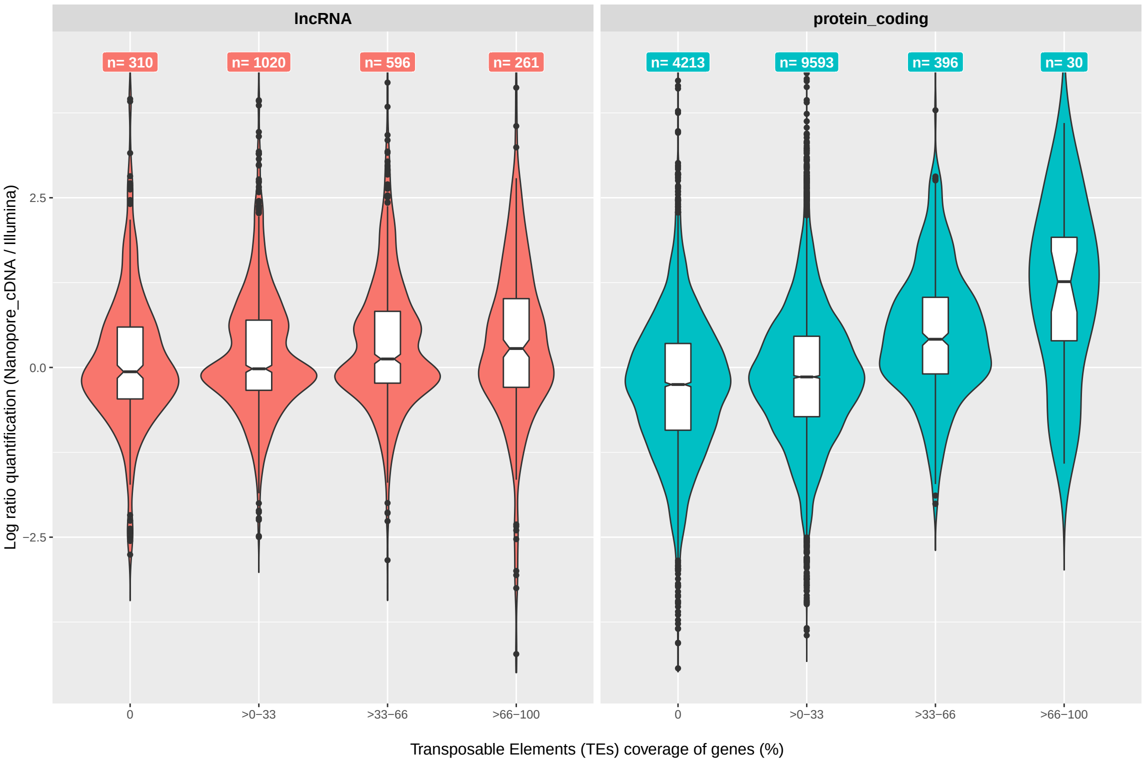


(b)

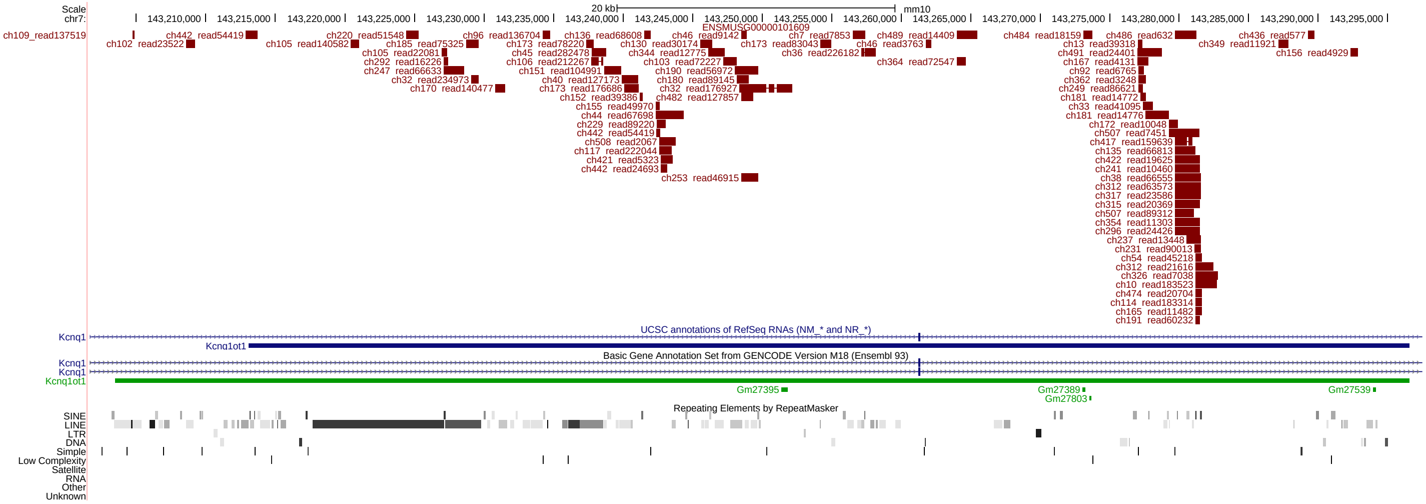


Figure 10: **Mouse lncRNA quantification in the ONT and Illumina cDNA-Seq.** (a) Gene quantification ratio (ONT cDNA-Seq versus Illumina cDNA-Seq) with variable gene coverage in Transposable Elements (TEs). Long non-coding RNAs (lncRNAs) are represented on the left panel (in red) while protein-coding genes are represented on the right (in blue). Only genes expressed in both conditions are represented (e.g. quantifications with $TPM > 0$ and $Nanopore_cDNA >= 1$). (b) UCSC screenshot of the KCNQ1OT1 locus. ONT cDNA-Seq reads mapped onto mm10 are represented on the top track, followed UCSC and GENCODE gene annotation tracks. Finally, the bottom tracks represents the repeat elements including transposable elements annotated by Repeat Masker.

(a)

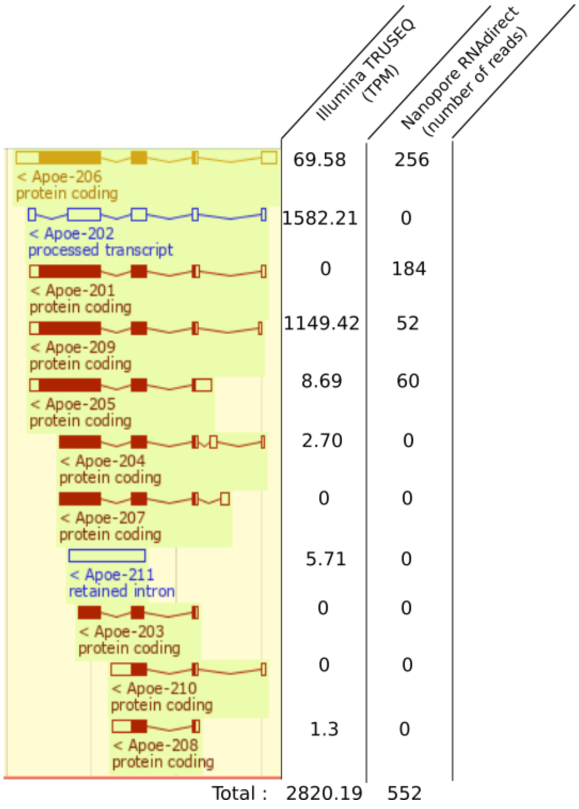


(b)

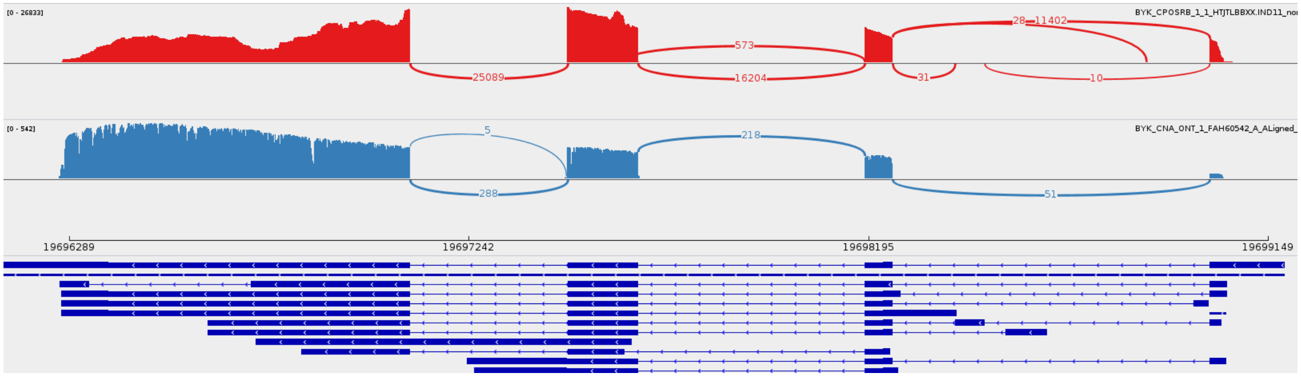


Figure 11: **Quantifications for APOE trancripts** (a) APOE annotation visualized with the Ensembl genome browser and quantification obtained with Illumina Truseq and RNAdirect (b) Sashimi plot obtained with IGV. Junctions covered by less than 5 reads were filtered out. First track shown Illumina Truseq reads and second track Nanopore RNA direct reads. As shown by the Sashimi plot, the most expressed transcript is not annotated. It Correspond to the transcript Apoe-206 with a shorter UTR.
